# High Throughput Fitness Profiling Reveals Loss Of GacS-GacA Regulation Improves Indigoidine Production In *Pseudomonas putida*

**DOI:** 10.1101/2021.02.02.429437

**Authors:** Thomas Eng, Deepanwita Banerjee, Andrew K. Lau, Emily Bowden, Robin A. Herbert, Jessica Trinh, Jan-Philip Prahl, Adam Deutschbauer, Deepti Tanjore, Aindrila Mukhopadhyay

**Affiliations:** Joint BioEnergy Institute, Lawrence Berkeley National Laboratory, Emeryville, CA 94608, United States; Biological Systems and Engineering Division, Lawrence Berkeley National Laboratory, Berkeley, California, USA; Advanced Biofuel and Bioproduct Process Development Unit, Lawrence Berkeley National Laboratory, Emeryville, CA 94608, United States; Environmental Genomics and Systems Biology Division, Lawrence Berkeley National Laboratory, Berkeley, California, USA

## Abstract

*Pseudomonas putida* KT2440 is an emerging industrial microbe amenable for use with renewable carbon streams including aromatics such as *para*-coumarate (*p*CA). We examined this microbe under common stirred-tank bioreactor parameters with quantitative fitness assays using a pooled transposon library containing nearly all (4,778) non-essential genes. Assessing differential fitness values by monitoring changes in mutant strain abundance over time identified 31 genes with improved fitness in multiple bioreactor-relevant parameters. Twenty-one genes from this subset were reconstructed, including GacA, a signaling protein, TtgB, an ABC transporter, and PP_0063, a lipid A acyltransferase. Twelve deletion strains with roles in varying cellular functions were evaluated for conversion of *p*CA, to a heterologous bioproduct, indigoidine. Several mutants, such as the Δ*gacA* strain improved both fitness in a bioreactor and showed an 8-fold improvement in indigoidine production (4.5 g/L, 0.29 g/g *p*CA, 23% MTY) from *p*CA as the carbon source.

## Introduction

Synthetic biology has the potential to produce many new molecules of interest which are challenging to synthesize by traditional chemistry. However, economical bioproduction at industrial scale depends on optimizing many parameters, including growth under bioreactor conditions, achieving high product titers, rates, and yields (TRY), as well as utilization of as many carbon streams derived from renewable carbon feedstocks. While many new molecules can be produced at the laboratory scale, successful development of an economically-viable strain at industrial scale (20,000 L - 2,000,000 L) is estimated to cost as much as 1 billion dollars^1^.

From an economics perspective, one of the most impactful ways to improve the viability of a process is to reduce the cost of the carbon biomass used as a substrate^2^. The use of lignocellulosic biomass in place of pure sugars as a low-cost feedstock could make these microbial processes financially feasible for high volume, low value molecules such as biofuels^3^. Currently, sugars are extracted from the cellulose and hemicellulose fractions, whereas the lignin fraction has proved to be challenging to convert biologically. Baral *et al* reported that in order to be cost-competitive with petroleum-derived jet fuels, bio-jet fuels would need to be sold at a market price around $2.5/gallon. Coproducts derived from lignin carbon streams are an underexplored avenue which could help satisfy this cost ceiling^3^, but lignin depolymerization can yield structurally distinct aromatic compounds, each of which could be used as a carbon source^4^. A solution from a recent report indicates that base-catalyzed lignin depolymerization could simplify this process, allowing for the recovery of a single dominant aromatic molecule, *para*-coumarate (*p*CA)^5^.

*Pseudomonas putida* KT2440 is a promising microbe with potential for biotechnology applications; first identified as a solvent and stress tolerant microbe, a spontaneous mutation in strain mt-2 improved plasmid transformation efficiencies^6,7^. *P. putida* KT2440 is able to grow using *p*CA as a sole carbon source^8^, giving it an advantage over other microbes like *Escherichia coli* or *Saccharomyces cerevisiae*. *P. putida* natively expresses Ttg-family efflux pumps to limit *p*CA toxicity, which may export the molecule and contribute to its tolerance of ~100 mM *p*CA^9–11^. *P. putida* has been recently engineered to convert aromatic compounds to heterologous metabolites^12,13^, but process validation for production in larger bioreactor formats is rare. For example, at the 300 L scale, production of a native compound, medium chain length polyhydroxyalkanoates (mcl-PHA) was optimized, but in a glucose feed regime^14^. Moreover, there are inherent differences in the conditions used to cultivate microbial strain in a shake flask vs. the conditions in which bioproduction will finally be deployed (e.g. uniform C source, pH, DO)^15,16^. This could be especially impactful on obligate aerobes such as *P. putida^17^*. To de-risk the scaling-up of any new microbial process, insights derived from cell physiology in stirred tank bioreactors could clarify how native cellular processes in *P. putida* are different from laboratory cultivation conditions.

While rationally-engineered gene deletions may improve specific aspects of cell growth or productivity, gene deletions in seemingly-unrelated processes have also yielded increases in heterologous protein activity^18,19^. These studies motivate the use of unbiased screens to identify factors which improve expression of heterologous gene products at bioreactor scales. Querying single gene mutants from a pooled *P. putida* mutant library could identify genes and regulatory networks required for robust growth in bioreactors. The quantitative fitness method using pooled barcoded transposon library we use is called RB-TnSeq and has been described for *P. putida* KT2440^20^.

In this work, we used fitness profiling data to identify candidates for reconstruction as isogenic deletion strains. In turn, we examined the bioconversion of *p*CA to a heterologous product, indigoidine, in bioreactors. We identified that inactivating a two component regulatory system (GacS-GacA, also known as UvrY-BarA in *E. coli*) led to improved product titers of the heterologous gene pathway for indigoidine production when fed *p*CA as a carbon source.

## Results

### A core cellular signature for growth under varied process parameters in bioreactors using functional genomics

We designed an experimental regime to identify *P. putida* mutants with changed fitness under conditions relevant to industrial cultivation. In contrast to lab scale experiments in shake flasks, culture tubes or microtiter plates, biomanufacturing processes for microbes implements cultivation in impeller-mixed jacketed tanks, where gases (ie, ambient air, oxygen) and nutrients (sugars, nitrogen sources) are added to the microbial culture during a given process^15,16,21^.

Using a pooled barcoded transposon mutant library in *P. putida* KT2440 we were able to rapidly evaluate ~100,000 unique transposon mutants covering nearly all (~4,800) non-essential genes with quantitative fitness assays. These cultures of pooled mutants were grown in bioreactors **(Figure 1A)** under conditions as outlined in **Table 1** to characterize differential fitness changes across timepoints and process conditions. Quantifying changes in barcode abundance allows rapid identification of the specific mutants and their respective fitness values in a workflow referred to as RB-TnSeq^22^. In *P. putida*, this method has been used in predicting carbon catabolic pathways and the characterization of growth inhibitors^23–25^. Comparing fitness values from stirred tank conditions to laboratory scale experiments would allow identification of mutants with fitness changes across format and process conditions to distinguish from mutations which generally impacted strain fitness across all conditions.

**Figure 1.**
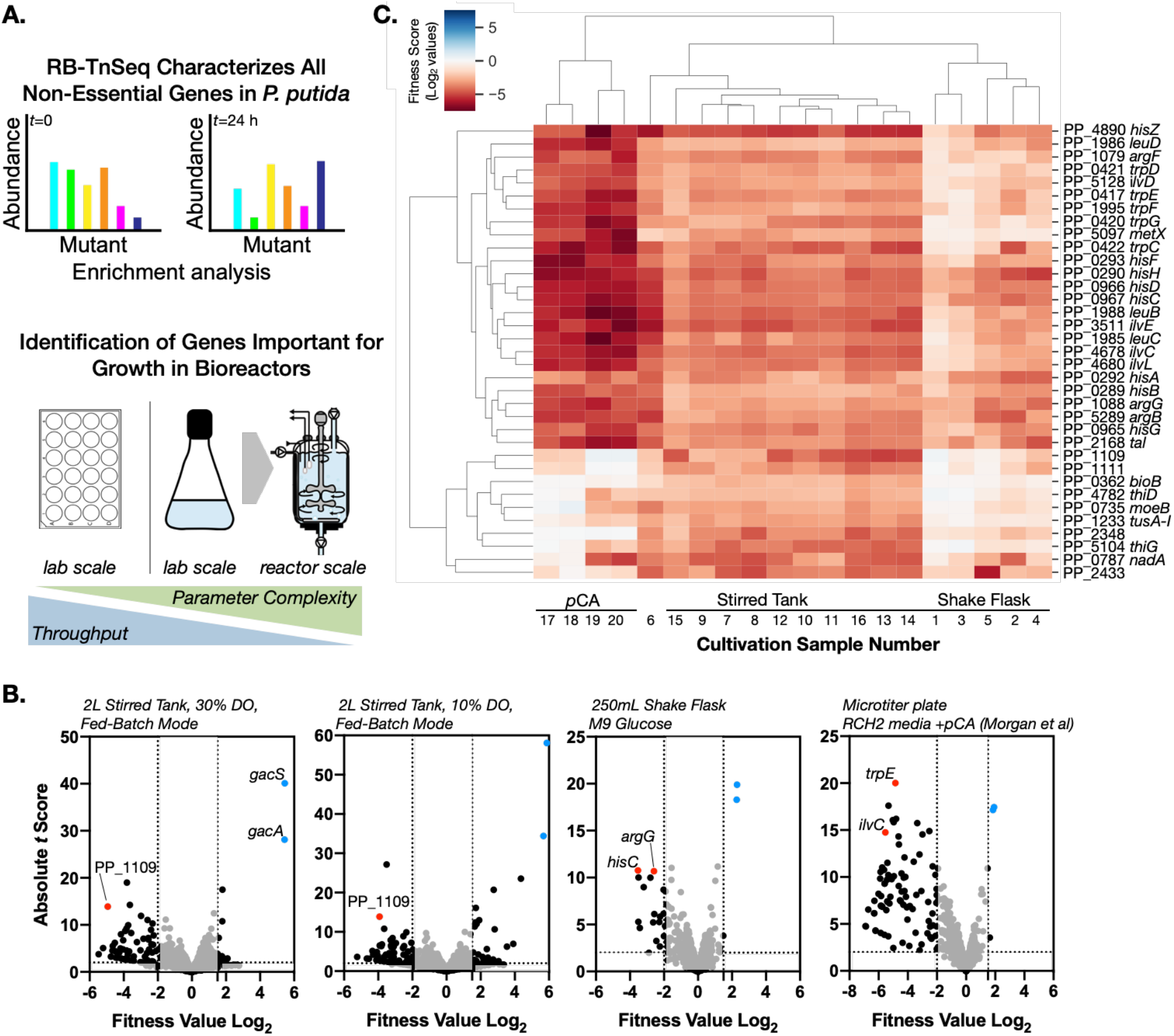
*Pseudomonas putida* Fitness Profiling Using Varied Process Parameters Conditions Reveals Gene Pathways Required for Robust Growth. a) Schematic of workflow. RB-TnSeq tracks differential mutant abundance across timepoints and conditions (refer to Table 1). Mutant abundance is tracked over time for a given condition and normalized to the initial abundance at T0. b) Volcano plots of four representative RB-TnSeq experiments. Strong fitness defects are indicated with dotted lines indicating cutoff values. For fitness, a cutoff threshold for log_2_ values > 1.5 or < −2.0 was used. For absolute *t* scores, the threshold chosen was *t* > 2. Fitness values for mutants in a two-component signaling system, GacS-GacA, is highlighted in blue. Several mutants that also appear in panel c are indicated in red. c) Hierarchical clustered heatmap for 35 gene mutants that were fitness-compromised for bioreactor conditions showing their corresponding fitness profile under laboratory cultivation with either glucose or *p*CA as the carbon source. Both genes and conditions are clustered.

**Table 1.** Summary of conditions and cultivation formats tested in this study for quantitative fitness analysis with the *P. putida* KT2440 RB-TnSeq library. All experiments were conducted at 30 °C. Samples 17 and 18 were described in Price *et al,* 2019^20^; samples 19 and 20 were described in Incha *et al* 2020^24^.

| No. | Sample ID | Culture Format/Scale | Base Media | Time point Sampled, Replicate Number | Feed Mode & Carbon Source | Dissolved Oxygen Setpoint |
| --- | --- | --- | --- | --- | --- | --- |
| 1 | Shake Flask-1 | 250 mL shake flask | M9 | 48h, R1 | Batch, Glucose | NA |
| 2 | Shake Flask-2 | 250 mL shake flask | M9 | 72h, R1 | Batch, Glucose | NA |
| 3 | Shake Flask-3 | 250 mL shake flask | M9 | 48h, R2 | Batch, Glucose | NA |
| 4 | Shake Flask-4 | 250 mL shake flask | M9 | 72h, R2 | Batch, Glucose | NA |
| 5 | Shake Flask-5 | 250 mL shake flask | M9 | 48h, R3 | Batch, Glucose | NA |
| 6 | Shake Flask-6 | 250 mL shake flask | M9 | 72h, R3 | Batch, Glucose | NA |
| 7 | 2L-1 | 2 L Sartorius Bioreactor | M9 | 24 h, R1 | Batch, Glucose | 10% |
| 8 | 2L-2 | 2 L Sartorius Bioreactor | M9 | 48 h, R1 | Batch, Glucose | 10% |
| 9 | 2L-3 | 2 L Sartorius Bioreactor | M9 | 72 h, R1 | Batch, Glucose | 10% |
| 10 | 2L-4 | 2 L Sartorius Bioreactor | M9 | 48 h, R2 | Batch, Glucose | 10% |
| 11 | 2L-5 | 2 L Sartorius Bioreactor | M9 | 72 h, R2 | Batch, Glucose | 10% |
| 12 | 2L-6 | 2 L Sartorius Bioreactor | M9 | 24 h, R1 | Batch, Glucose | 30% |
| 13 | 2L-7 | 2 L Sartorius Bioreactor | M9 | 48 h, R1 | Batch, Glucose | 30% |
| 14 | 2L-8 | 2 L Sartorius Bioreactor | M9 | 72 h, R1 | Batch, Glucose | 30% |
| 15 | 2L-12 | 2 L Sartorius Bioreactor | M9 | 24 h, R1 | Fed Batch, Glucose | 30% |
| 16 | 2L-13 | 2 L Sartorius Bioreactor | M9 | 40 h, R1 | Fed Batch, Glucose | 30% |
| 17 | *pCA*-1 | 24 well plate | RCH2 | R1 | Batch, *pCA* | NA |
| 18 | *pCA*-2 | 24 well plate | RCH2 | R2 | Batch, *pCA* | NA |
| 19 | *pCA*-3 | 96 deep well plate | MOPS | R1 | Batch, *pCA* | NA |
| 20 | *pCA*-4 | 96 deep well plate | MOPS | R2 | Batch, *pCA* | NA |

For each sample, we were able to calculate the fitness and a corresponding *t*-score for single transposon mutants for each of the 4,778 genes in the pooled library. The calculated fitness value for each timepoint is the log_2_ ratio of the population abundance for the sampled timepoint over the initial mutant abundance measured at the start of each experimental time course. The *t-* score assesses how reliably a fitness value is different from zero. For most conditions, most genes do not have a measurable differential fitness value and therefore have fitness values and *t* scores close to 0. For our genome-wide analysis, we selected strong, statistically significant determinants of fitness and demanded that fitness values must be >1.5 or <-2 with an absolute *t* score of >2. Volcano plots of mutant fitness values and their corresponding *t* scores are plotted for five representative experiments in **Figure 1B**.

From this dataset we identified thirty-five transposon mutants in *P. putida* which displayed growth defects under these conditions as displayed in **Figure 1C**. Hierarchical clustering of mutants that had decreased fitness (less than −2) indicated that most bioreactor samples fed glucose were similar to samples fed glucose in the shake flask format, but had different responses when cells were fed *p*CA. For conditions listed in **Table 1**, mutants with decreased fitness were recovered in near-complete amino acid biosynthesis pathways for methionine, and tryptophan. Other amino acid pathways were partially identified, such as for leucine, arginine, and aspartate. Pathways predicted for sulfur relay and thiamine biosynthesis (PP_0261, PP_1233, PP_5104) or metal ion homeostasis (PP_3506, PP_0910) were also implicated for robust growth under bioreactor conditions (**Supplementary Data 1**). We observed that when the genes were clustered, two additional uncharacterized genes (PP_0292, PP_0289) also were present in this group, suggesting they have related functions. Nineteen mutants were unique to growth on *p*CA. These included transcriptional regulators and metabolic genes in pathways related to aromatic compound catabolism already described elsewhere^24^. Including *p*CA fitness profiling data from the Price *et al^20^* dataset also strengthened evidence for statistically significant fitness defects in other metabolic genes including PP_5095/*proI* (involved in proline biosynthesis), PP_0356/*glcB* (malate synthase), PP_4700/*panC* (pantothenate synthase) and PP_4799 (a putative muramoyltetrapeptide carboxypeptidase).

A number of regulatory systems were also identified that have not been found to have a fitness phenotype in earlier studies, suggesting that bioreactor conditions generated several previously uncharacterized global stress responses. Deletion of either the sigma-38 stress response sigma factor (PP_1623/*rpoS*) or housekeeping sigma factor sigma-70 (PP_4208) strongly decreased fitness in the bioreactor. Other *P. putida* RB-TnSeq datasets did not show the deletion of either sigma factor to have fitness defects (unpublished results, RB-TnSeq fitness browser, https://bit.ly/3bifz2h), suggesting that the fitness enhancement in a bioreactor is enhanced by both sigma-38 and sigma-70 transcriptional regulation, and that their regulatory network is not redundant. Additionally, four environmental/nutrient availability sensing two-component signaling systems were also implicated in bioreactor fitness **(Figure 1, Supplementary Data 1)**: a nitrogen stress sensor (PP_2388-PP_2390)^26^; a sensor implicated in chloramphenicol resistance (PP_0185)^27^; a sensor implicated in lipid A remodeling (PP_2348); and a two component system important for adaptation to growth in minimal media (PP_4505-PP_2714). The specific environmental signals which activate these remaining two component signaling systems have not been identified.

Other mutants which decrease fitness in bioreactors included mutations in PP_5303/*ridA*, a reactive oxygen species responsive chaperone, or PP_0735/*moeB*, an adenyltransferase which adenylates molybdopterin synthase, were deficient for growth. Finally, mutations in four other genes could not be assigned a function due to low homology to previously characterized genes or correlation with known processes. In summary, we have identified a core cellular signature for growth under a variety of common process parameters in bioreactors using a functional genomics approach.

While the negative fitness values from the RB-Tnseq method identified cellular sensitivities in bioreactors, we reasoned that mutants with improved fitness values could also be leveraged for biotechnological applications. Hierarchical clustering of positive fitness mutants across all bioreactors in comparison with *p*CA and shake flask conditions indicated that a fitness signature in the bioreactor was distinct from either standard laboratory format using the aromatic carbon source or glucose (**Figure 2A**). We identified transposon mutants in eighteen genes which consistently exhibited quantitatively improved fitness under these growth conditions in a bioreactor and are plotted with hierarchical clustering in **Figure 2A**. Additionally, a two component signaling system, *gacS-gacA* (PP_1650-PP_4099), was routinely recovered as a fitness enhanced mutant under many conditions (**Figure 2A** and **2B**).

**Figure 2.**
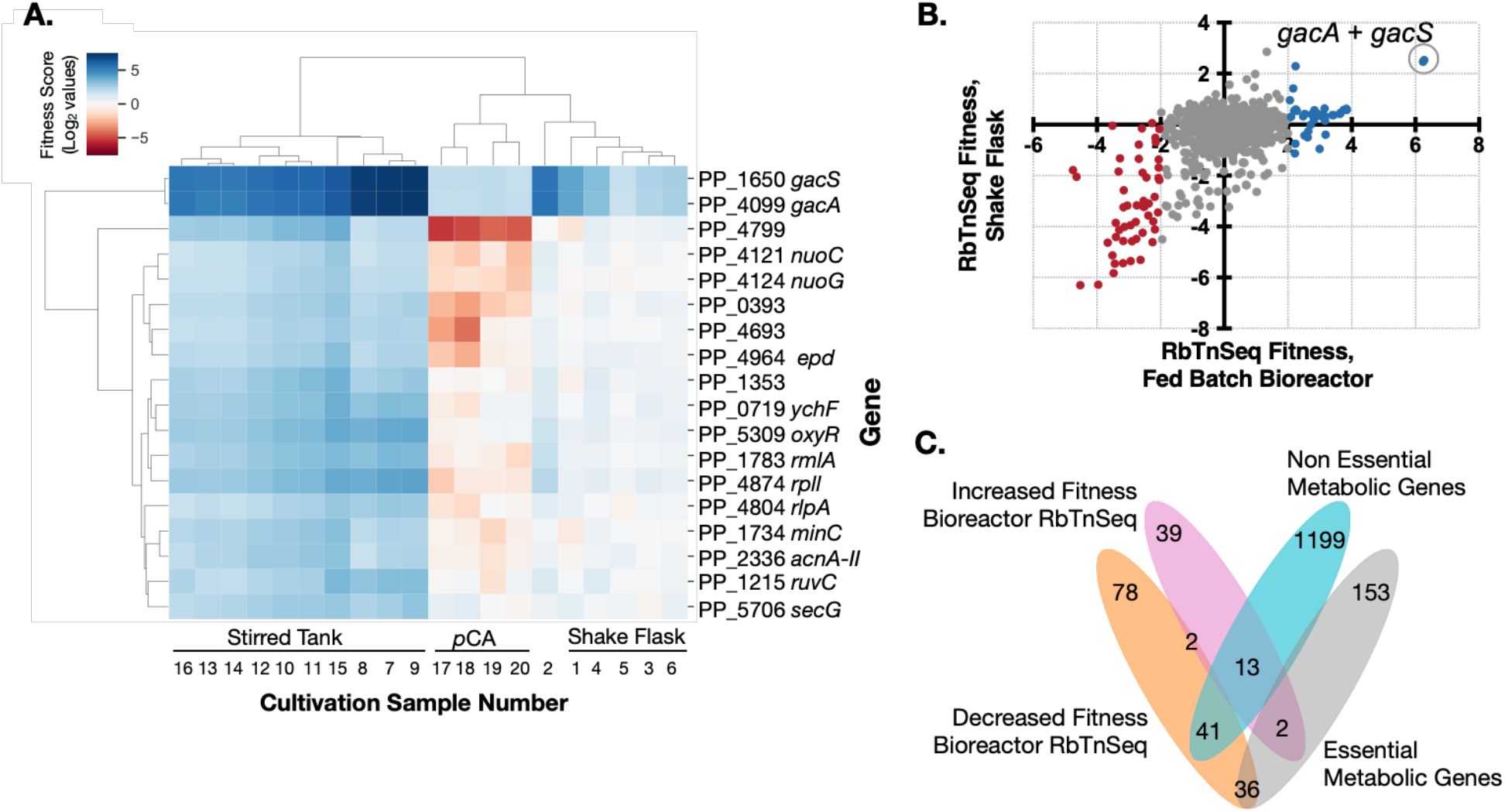
Identification of *P. putida* Enhanced Growth Mutants. a) Heatmap for 18 gene mutants with enhanced fitness (fitness value greater than 1.5) across all bioreactor condition data points compared to fitness value under laboratory conditions using *p*CA in microtiter plates or glucose in shake flask as the sole carbon source. Refer to Table 1 for a full description of conditions corresponding to the sample numbers. b) Scatter plot showing the fitness values of mutants with enhanced fitness (red, fitness values >2) or compromised fitness (blue, fitness values <2) under grown in shake flask *vs.* a fed-batch bioreactor. c) Venn diagram indicating distribution of genes binned into four different categories from the transposon mutant pool using fitness profiling values from bioreactor fitness experiments and gene essentiality.

Approximately half of the mutants identified from these fitness profiling experiments were related to metabolic processes (**Figure 2C**). The remaining non-metabolic candidates (described in **Supplementary Data 2**) encoded a diverse range of cellular functions, such as PP_1215/*ruvC* (a crossover junction endodeoxyribonuclease), PP_1353 (an uncharacterized conserved membrane protein), and PP_5309 (a LysR-family transcriptional regulator). Inactivating PP_1428/*rpoE* (Sigma factor sigma-E) led to a slight fitness improvement in most of the bioreactor conditions tested by RB-TnSeq, but not in the control shake flask experiments. Many of these genes likely encode global master regulators and their deletion have pleiotropic impacts across cell physiology. In summary, we identify these gene loci that are potential gene targets (**Table 2**), whose inactivation would result in improved fitness in a bioreactor including metabolic genes as well as non-metabolic global regulators across varied oxygen and mixing conditions in bioreactors.

**Table 2.**
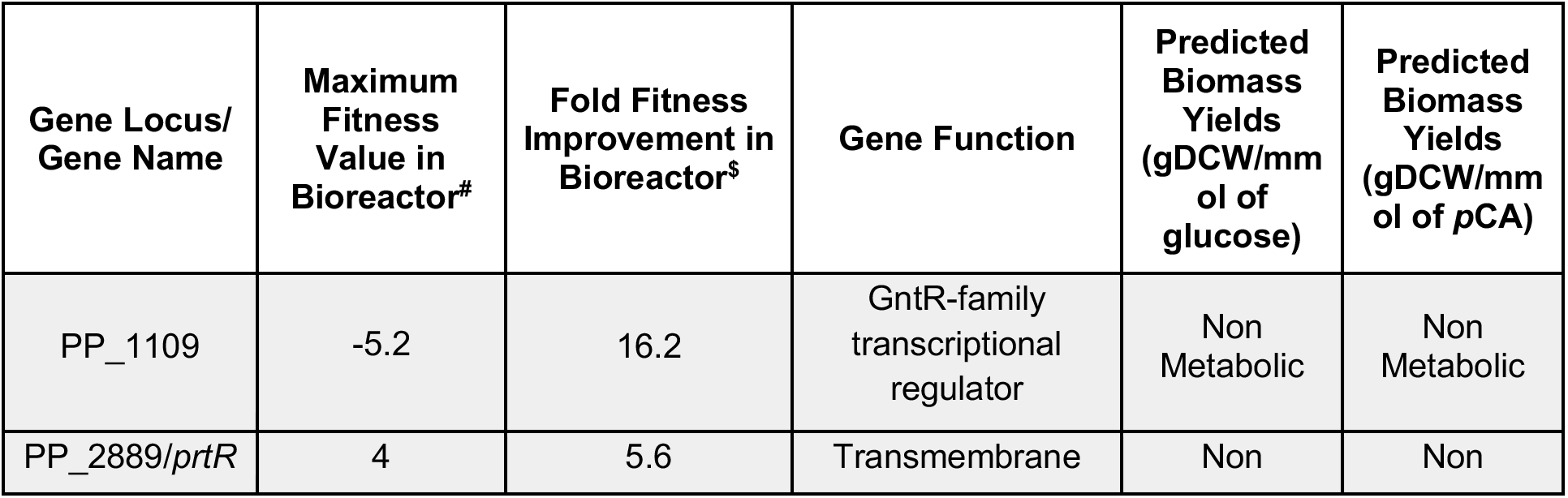
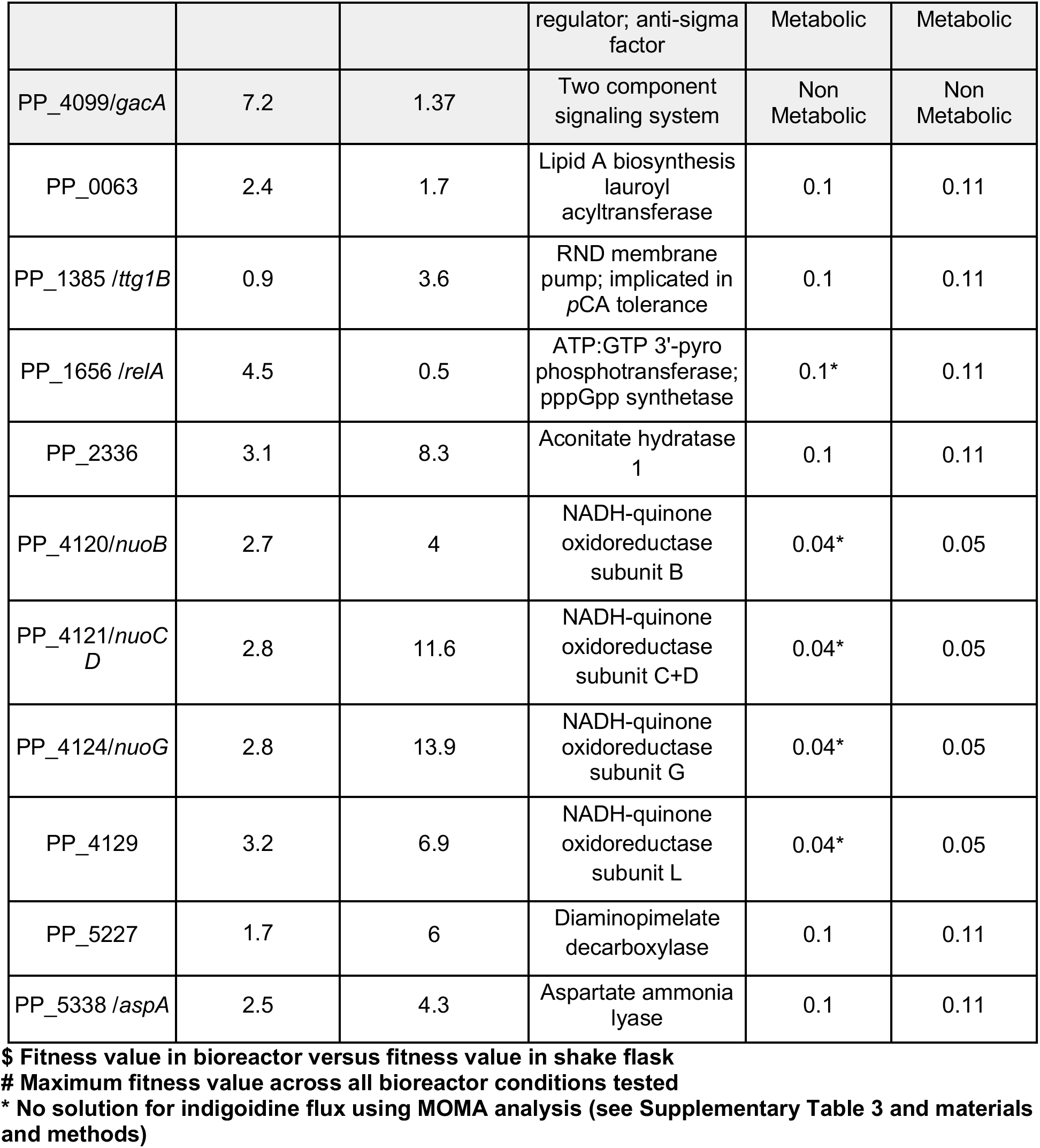
Fitness-Enhanced Deletion Mutants in this Study. Genes are first sorted as metabolic or non-metabolic, and next sorted according to genomic locus ID from smallest to largest values. If known, common gene names are also indicated. The complete list of loci targeted for deletion, including newly identified essential genes, is described in Supplemental Table 1.

### The bioconversion of a lignin derived aromatic to a heterologous bioproduct

The RB-TnSeq functional genomics analysis indicated that inactivating a small number of cellular processes would lead to improved fitness in a bioreactor. In addition, we included two other analyses to capture additional useful improvements. Specifically, we included mutants which have higher fitness values in stirred tank bioreactors when compared to the shake flask format. We calculated differential fitness values as the ratio of log_2_ fitness values in a bioreactor over shake flask cultivation as the denominator, which indicated a small number of genes should be included even though they did not have strong absolute fitness improvements or *t* scores that would otherwise meet the threshold. From this analysis 14 additional genes related to metabolism were identified. We evaluated how deleting these individual metabolic genes for their potential impact on maximum biomass yields using minimization of metabolic adjustment (MOMA) method^28^ when fed glucose or *p*CA as carbon sources. MOMA analysis predicted the immediate effect of a gene deletion with minimum perturbation in the metabolic flux distribution compared to wild type *P. putida*. Of the 14 genes, PP_0290 was predicted to be essential *in silico* for growth using both glucose or *p*CA as sole carbon source. All thirty-three genes that met at least one of these criteria are described in **Supplementary Table 1.**

The RB-TnSeq workflow under these process conditions enabled rapid characterization of nearly all non-essential *P. putida* mutants (**Figure 1A**). However, to use these improved chassis we built isogenic deletion mutants for each enhanced fitness mutant using allelic exchange plasmids targeting each locus for deletion **(Figure 3A and Supplementary Table 1)**. Consistent with the potential for heterozygous alleles in *P. putida^29^*, we were unable to generate deletion mutants for twelve candidate genes **(Supplementary Table 1),** but were successful in completing a library of thirteen deletion mutant strains (**Table 2**) to test for heterologous bioproduct formation, as modeled with the 2 gene non-ribosomal peptide (NRP), indigoidine. Indigoidine is generated from the condensation of two glutamine molecules (**Supplementary Figure 1**) and is catalyzed by a heterologous non-ribosomal peptide synthetase (NRPS) based pathway^30–32^.

**Figure 3.**
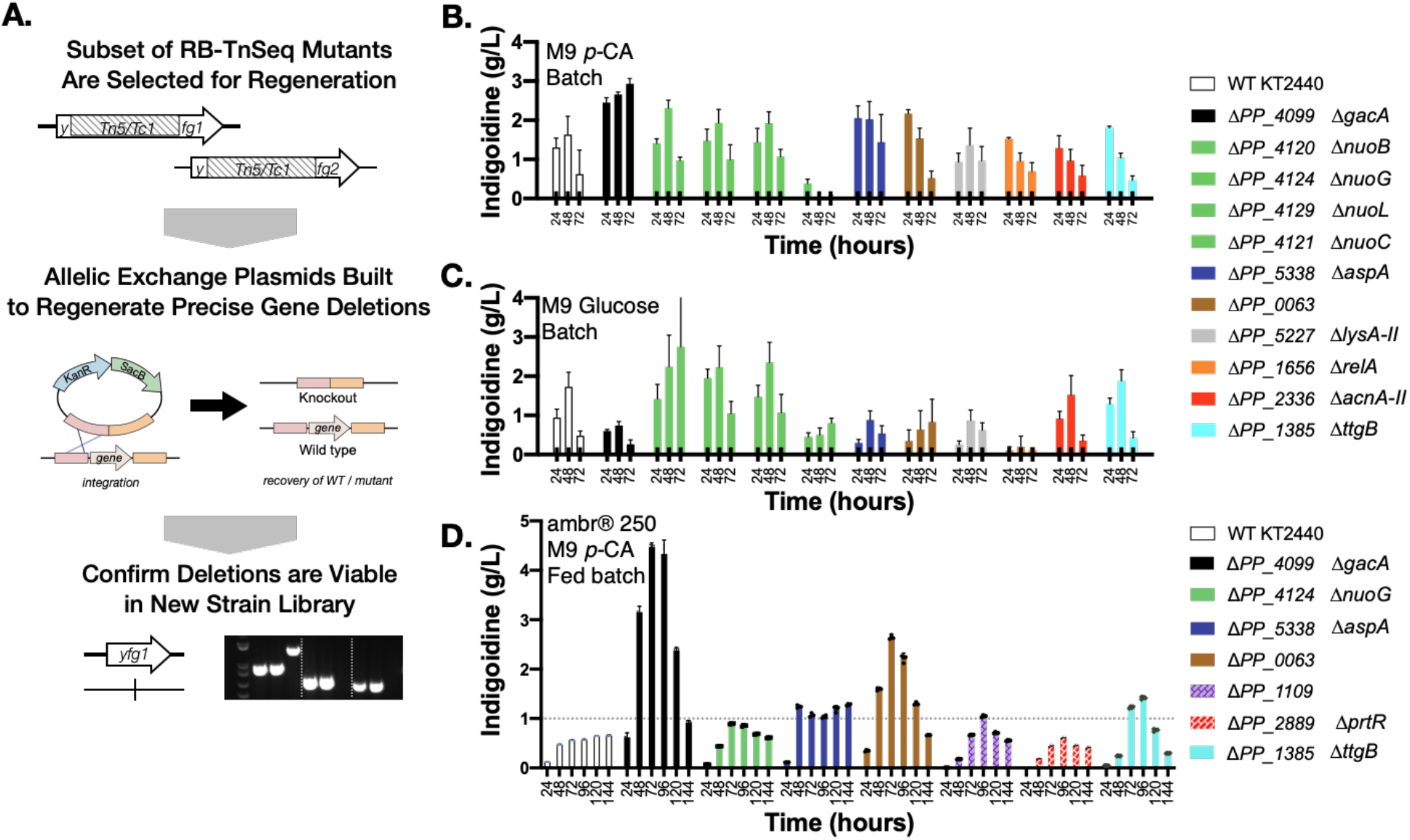
Indigoidine productivity in isogenic deletion strains across scales. A. Workflow to build new isogenic deletion mutants. B. Indigoidine production in batch mode using *p*CA as carbon source C. Indigoidine production in batch mode using glucose as the carbon source. Single deletions in the nuo holocomplex are indicated in green. Otherwise, deletions are arranged by decreasing titer. For B and C, error bars indicate SD and n=3 from independent biological replicates. D. Fed-batch mode production of indigoidine using *p*CA as the carbon source from n=3 technical replicates.

To test indigoidine production, we selected a subset of mutants that showed the most promise either by metabolic modeling or by mining the literature for potential roles in processes related to glutamine synthesis, the immediate precursor to indigoidine. The benefit to biomass formation ideally should not come at the cost of bioproduct titer, rates, and yields^33,34^. Deletions of metabolic genes were also analyzed for their potential impact on indigoidine titer. Using a genome scale metabolic model of *P. putida*, iJN1462^35^ and Flux Balance Analysis (FBA), we calculated maximum theoretical yields (MTY) of indigoidine and its precursors for this carbon/substrate pair (**Supplementary Table 2**). This carbon source to final product MTY pair (*p*CA/indigoidine) of 0.66 mol/mol is higher than MTY calculated for the glucose/indigoidine pair of 0.54 mol/mol^36^. The predicted flux towards indigoidine in these mutants is summarized in **Table 2** and **Supplementary Table 3**. For several mutants, indigoidine yields were unlikely to substantially improve yield, but would still allow yields approximately 80 - 100% of WT. Six of the thirteen deletion mutants analyzed had no solution when calculating indigoidine flux using MOMA analysis when fed glucose, but solutions did exist for *p*CA feed conditions **(Table 2)**. These model predictions suggested there might be improvements to final product titer in these deletion strains. The indigoidine production pathway was then integrated into these deletion strains to produce the heterologous final product.

We selected thirteen deletion mutants for indigoidine production in the laboratory scale format which had various range of biological functions, fitness values, or biomass predictions from MOMA analysis (**Table 2**). These included 9 metabolic genes and Δ*gacA* among the remaining non metabolic genes. As controls, we included Δ*ttgB* to test if reducing *p*CA efflux could allow greater substrate availability for catabolism^27,37^; ΔPP_1109 which exhibited negative fitness values in the bioreactor; and ΔPP_2889 which the RB-TnSeq data was more fit only under batch-mode conditions but not fed-batch modes **(Table 1)**. The remaining genes in **Table 2** showed either differential or absolute fitness improvements in the bioreactor scale compared to the shake flask format.

Strains were assayed first in 24-deepwell plates to compare indigoidine production using either glucose or *p*CA as the carbon source. In this format, the WT strain produced about 1.5 g/L of indigoidine from either glucose or *p*CA as the carbon source after 48 hours of cultivation. In contrast, Δ*gacA* strains produced 2.5 g/L of indigoidine after 48 hours using *p*CA (**Figure 3B**) but only 0.5 g/L indigoidine from glucose (**Figure 3C**). Several subunits of the NADH-quinone oxidoreductase complex (PP_4120, PP_4124, PP_4129) led slight improvements in indigoidine titer when cells were fed glucose, but not *p*CA. While these proteins are thought to function as part of a single holocomplex, the differences in indigoidine production are consistent with the differences in transposon mutant fitness **(Table 2)**. Deletion strains ΔPP_2889, ΔPP_5338, ΔPP_0063, ΔPP_5227, ΔPP_1656, ΔPP_2336, and ΔPP_1385 also showed improved indigoidine titer on *p*CA but not on glucose **(Figure 3, Supplementary Figure 2)**. ΔPP_1109 which had a negative fitness in bioreactors, did not improve indigoidine titer **(Supplementary Figure 2).** In summary, we identified several mutants with improved indigoidine production from *p*CA, which allowed us to further down-select candidate strains for bioreactor runs.

Automation-assisted fed-batch bioreactors (Ambr^®^ 250) enable medium throughput analysis in stirred tank bioreactors and were used to examine the most promising four deletion mutant strains for indigoidine production (Δ*gacA*, ΔPP_4124, ΔPP_5338, and ΔPP_0063). In this scaleup, Δ*gacA* strains produced 4.5 g/L indigoidine after 72 hours, whereas the WT strain produced 0.5 g/L in the same timeframe. ΔPP_0063 also showed some improvement over the WT strain with a titer of 2.5 g/L. Deletion strains ΔPP_4124 or ΔPP_5338 did not further improve indigoidine titer in the bioreactor. The remaining control strains performed as expected; a representative deletion strain with a negative fitness value (ΔPP_1109) did not produce more indigoidine than wild type; reducing *p*CA efflux (Δ*ttgB*) or optimizing for growth under batch mode conditions (ΔPP_2889) also failed to improve titer. The indigoidine yield from the control strain was 0.034 g indigoidine / g *p*CA, and the yield from the Δ*gacA* production strain was 0.29 g indigoidine/ g *p*CA, an 8.5 fold improvement over wild type. The Δ*gacA* strain reached 29% MTY (indigoidine / *p*CA) under fed-batch conditions. This result demonstrates a successful application of fitness profiling of deletion libraries for improved bioconversion route to produce indigoidine when fed a lignin-derived monomer as the sole carbon source.

## Discussion

In this study, we have used high throughput functional genomic approaches to identify genetic enhancers in *P. putida* which improve growth in bioreactors under a range of process conditions. *P. putida* is of special interest as a host for scalable and sustainable production due to its ability to metabolize aromatic components of plant feedstocks such as *p*CA. The key discovery in this study is that we identified a gene target that improves fitness in a bioreactor and also under certain conditions enhances production of a model heterologous product. Our study describes the distinct fitness landscape of scaleup conditions on cells, explaining why larger format processes are so unpredictable based on performance at the laboratory scale. Due to the pooled nature of our high-throughput assay, disrupted gene pathways that utilize metabolites which can be complemented by secreted metabolites from other mutants in the population will not be detected in this assay. Regardless, our negative fitness mutants identify important genes to avoid inactivating when considering genome scale approaches for host optimization^38^.

We have quantified the differences between bioreactors and shake flasks and demonstrated that stirred tank bioreactor conditions pose a burden on cell physiology with a distinct signature from that of conventional laboratory growth conditions in the same minimal media. An earlier report examined the budding yeast deletion collection and observed that analagous defects in amino acid biosynthesis pathways impaired cell survival under low temperature and high pressure conditions^39^. We also show that global stress response pathways, such as sigma-38 (RpoS) and sigma-70 (RpoD), were implicated as providing non-redundant stress responses for efficient growth in bioreactors. These stress-responsive transcription factors can parse many different nutritional and environmental changes; for example, in *E. coli* the RpoS response is modulated by intracellular glutamate concentrations. Changes in glutamate binding to the RpoS holoenzyme dictate the subset of activated downstream genes in a stress response^40^. While bioreactor conditions may be far more controlled compared to its native soil habitat, this study indicates that *P. putida* has a full genetic complement ready to respond to the heterogeneous oxygen and variable nutrient cycles we insult these cells with.

Scalability is unpredictable; not all global regulators are required for robust growth in stirred tank bioreactors. In a related study, the Crc protein was identified as a global regulator whose inactivation has been demonstrated to improve product formation (cis, cis muconic acid) from *p*CA under laboratory cultivation conditions^13^. However, when translated to an aromatics and sugar co-utilizing engineered *P. putida* strain, *crc* deficient cells exhibited a significant lag phase under bioreactor growth conditions^41^. As we did not identify *crc* as a positive fitness mutant in our analysis of bioreactor-advantaged strains, we instead argue optimizing strains for bioproduct formation using laboratory settings may be inadequate; the same optimizations can have negative implications upon scale-up.

In *Pseudomonas aeruginosa*, the GacS-GacA system is known to be implicated in a remarkably wide range of conditions^42–44^ and in *P. putida,* was recently shown to have a role in muconate production from glucose^45^. In our study, the GacS-GacA two component signaling system was found to not only be a prominent deletion target for improving fitness but also an unanticipated route to improve heterologous product formation from *p*CA. A meta-analysis of functional genomics data from all public RB-TnSeq datasets (**Supplementary Table 4 and 5**) indicates that the *P. putida* GacS-GacA system may have little crosstalk amongst other signaling systems, as mutants in *gacS* strongly phenocopy *gacA* mutants (**Figure 4A**). In contrast, a signaling correlation analysis of the better characterized *E. coli* homologous GacS-362 GacS-362 GacA system shows a weaker correlation (r^2^ for *P. putida*=0.997; r^2^ for *P. stutzeri*=0.995; r^2^ for *E. coli*=0.688). Moreover, the absolute fitness values for *E. coli gacA* or *gacS* homolog mutants do not indicate any strong, biologically relevant phenotypes. With only ~25% identity between *E. coli* and *P. putida gacS* homologs, knowledge from *E. coli* is not translatable to *P. putida*. Regardless, this dataset implies that there may be important differences in how these signaling systems function between these organisms, which biases our literature survey to favor experimental evidence from *Pseudomonads* over *E. coli*.

**Figure 4.**
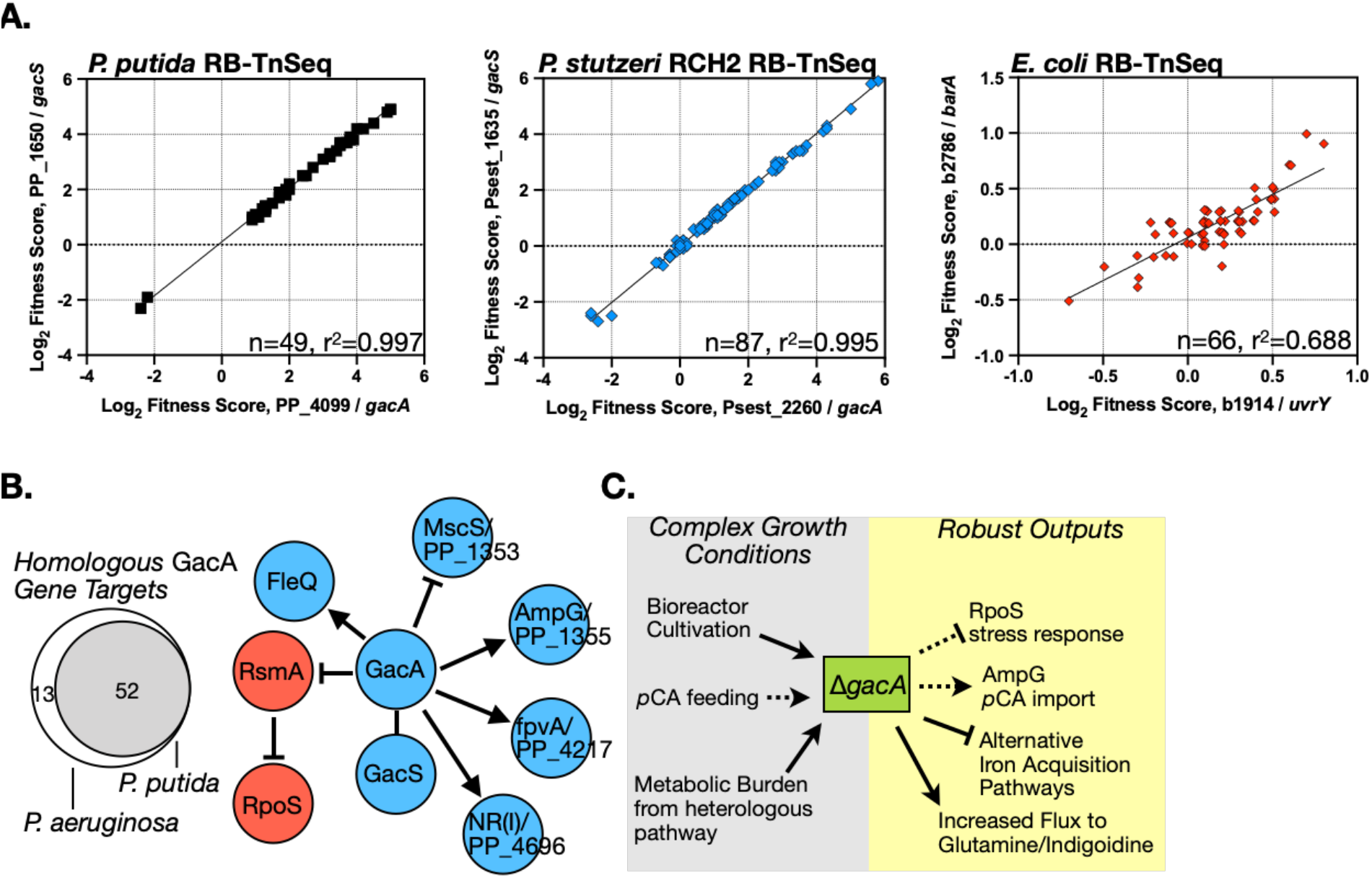
Functional Differences Between GacS-GacA And Putative *E. coli* Homolog BarA-UvrY. A. Meta-analysis of fitness values for GacS-GacA mutants across several different conditions for three different microbes, *P. putida*, *P. stutzeri* and *E. coli* using RB-TnSeq data. B. and C. Model. Δ*gacA* cells parse many nutrient and environmental signals and improve formation of a glutamine-derived product, indigoidine.

The deletion libraries chosen to test for improved indigoidine production profiling based on the fitness landscape of *P. putida* under bioreactor conditions had different production phenotypes when using these two carbon sources, specifically the Δ*gacA* deletion strain. Glucose is consumed through the ED-EMP pathway^46^ whereas *p*CA is utilized through the beta ketoadipate pathway^10^. Several known downstream targets of GacA (small RNAs rsmX, rsmY, rsmZ; grxD)^47,48^ are not included in the RB-TnSeq analysis pipeline, which is a shortcoming of this method for identifying small RNA based regulation. However, a stringDB meta-analysis^49^ also identified potentially conserved protein-protein interactions between GacA and other relevant candidate effector proteins **(Supplementary Figure 3)**. We hypothesize that an active GacS-GacA signalling system may induce formation of diverse secondary metabolites, biofilm formation and alternative iron sequestration pathways and these pleiotropic processes limit the carbon flux available for indigoidine production **(Figure 4B)**. In parallel, the constitutive derepression of an outer membrane permease (PP_1355), also regulated by this signaling system, could improve *para*-coumarate transport. Consistent with this, the inactivation of PP_1355 caused a fitness defect when cells were grown on *p*CA as a carbon source^20^. The improvement to indigoidine titer in Δ*gacA*, Δ*ttgB and* ΔPP_0063 strains, the three best indigoidine producers, occurred with *p*CA rather than glucose. While additional experiments are required to fully understand this outcome, aromatic molecules like toluene are known to induce a starvation response^50^ and occur in *p*CA. Without *gacS* present, the *rpoS* response is dampened^51^. This model is supported with our experimental data, as inactivation of *gacS* improves *P. putida* fitness when cells are grown on *p*CA. Although Δ*ttgB* had a slight fitness improvement under bioreactor conditions (**Table 2**), the indigoidine improvements were not as high as with the Δ*gacA* strain (**Figure 3B and 3D**). PP_0063 has been reported to play a role in the *P. putida* global stress response when cells were fed benzoate, a similar aromatic compound^52^. ΔPP_0063 strain had improved indigoidine titers compared to wild type but lesser than Δ*gacA* strain, suggesting that inactivating the PP_0063 regulatory network is not as beneficial towards indigoidine production as a Δ*gacA* deletion. In summary, our data suggests that final indigoidine titer is improved in the Δ*gacA* strain because a subset of starvation response genes are induced by *p*CA without activating the full complement of GacS-GacA regulatory targets. It is the context of *p*CA vs glucose cultivation that reveals the indigoidine productivity benefit. We speculate that a renewed focus on regulatory networks in this microbe will lead to improved optimization strategies for robust growth under dynamic environmental conditions with non-native carbon streams.

Our study advances the field of host engineering for heterologous bioproducts by applying methods in functional genomics to characterize host physiology under industrially relevant bioreactor conditions. Beyond providing a valuable new *P. putida* strain that converts the plant-derived aromatic *p*CA to the NRP indigoidine, this study provides a robust workflow to downselect strains for examination in the lower throughput but higher scale ambr^®^ 250 or bioreactor systems. The ambr*^®^* 250 improves our throughput to up to 12 simultaneous stirred tank runs but increasing the throughput above 24 units requires a significant capital investment. The isogenic deletion strain collection of bioreactor-advantaged mutants is ready to be screened with other heterologous gene pathways and carbon streams, such as xylose^41^. This functional genomics data can also help improve predictability of machine learning tools, such as ART^53^. Alternatively, the pooled library could be expanded to include double mutants or over-expressed genes to identify additional mechanisms of improved bioreactor growth. These strategies have the potential to identify better suited microbes for use in the emerging bioeconomy.

## Methods

### Chemicals, media and culture conditions

All chemicals and reagents were purchased from Sigma-Aldrich (St. Louis, MO) unless mentioned otherwise. When cells were cultivated in a microtiter dish format, plates were sealed with a gas-permeable film (Breathe-easy Sealing membrane, Sigma-Aldrich, St. Louis, MO). Tryptone and yeast extract were purchased from BD Biosciences (Franklin Lakes, NJ). Engineered strains were grown on M9 Minimal Media (NREL Formulation)^11^ with 10 g/L *para*-coumarate at 200 rpm at 30 °C. Overnight cultures of *P. putida* were grown and adapted in 5 mL M9 Minimal Media from single colonies. After three sequential rounds of adaptation, these cultures were used to inoculate cultures for indigoidine production runs at a starting OD_600_ of 0.1. All experiments were performed in triplicates and in different production scales. These included 200 μl culture volume in 24-deepwell plates (Axygen Scientific, Union City, CA), 2 mL culture volume in (InFors Multitron HT Double Stack Incubator Shaker), 999 rpm linear shaker, 70% humidity and 60 mL culture volume in 250 mL Erlenmeyer shake flask, 200 rpm orbital shaker). 0.3% w/v (20 mM) L-arabinose was used for (indigoidine production) induction of *bpsA-sfp* genes under the pBAD promoter.

### Strains and strain construction

*Pseudomonas putida KT2440* was used as the host for strain engineering. Electroporation with the respective plasmid (**Supplementary Table 6**) was performed using a BioRad MicroPulser preprogrammed EC2 setting (0.2 cm cuvettes with 100 μL cells, ~5 msec pulse and 2.5 kV). Transformed cells were allowed to recover at 25 °C for around 2.5 hours followed by plating onto selective media (containing respective antibiotics) followed by overnight incubation at 30 °C. Positive clones were confirmed by genotyping the respective locus by colony PCR using Q5 Polymerase (NEB, Ipswitch, MA) as described in the next section.

### Generation of Isogenic Deletion Strain Library

Open reading frames (ORFs) targeted for gene deletion were identified from the RB-TnSeq fitness values. Allelic exchange plasmids were constructed using either backbone pEX18GM or pK18mobsacB as previously described^11^. All homology arms generated for allelic exchange were sequence verified with Sanger sequencing (Genewiz Technologies, Waltham, MA). Deletions in *P. putida* KT2440 were generated as described in Mohamed *et al^11^* using 50μg/mL kanamycin or 30 μg/mL and gentamicin and subsequent counterselection on solid agar media supplemented with LB broth and 10% (w/v) sucrose. Kanamycin sensitive, sucrose resistant clones were then verified for the loss of the wild type locus using colony PCR primers flanking the targeted genomic locus using colony PCR. All primer sequences and allelic exchange plasmids are available post-publication from public-registry.jbei.org. All strains used in this study are provided in **Supplementary Table 7**.

### Analytics/ Sugar Quantification - HPLC

Glucose, *p*CA, and organic acids from cell cultures were measured by an 1100 Series HPLC system equipped with a 1200 Series refractive index detector (RID) (Agilent) and a Diode array detector (DAD) along with Aminex HPX-87H ion-exclusion column (300 mm length, 7.8 mm internal diameter; Bio-Rad Laboratories, Inc., Hercules, CA). 300 μL aliquots of cell cultures were removed at the indicated time points during production and filtered through a spin-cartridge with a 0.45-μm nylon membrane, and 10 μL of the filtrate was eluted through the column at 50°C with 4 mM sulfuric acid at a flow rate of 600 μL/min for 30 min. Metabolites were quantified by using external standard calibration with authentic standards.

### Indigoidine Quantification

Briefly, pelleted 100 μL of cells by centrifugation at 15000 rpm for 2 min. The supernatant was discarded and 500 μL DMSO was added to the pellet. The solution was vortexed vigorously for 30 s to dissolve Indigoidine. After centrifugation at 15000 rpm for 2 min, 100 μL of DMSO extracted indigoidine was added to 96-well flat-bottomed microplates (Corning Life Science Products, Corning, NY). Indigoidine was quantitated by measuring the optical density at 612 nm wavelength (OD612) using a microplate reader (Molecular Devices Spectramax M2E) preheated to 25 °C. Accounting for the any dilution applied, indigoidine was quantitated using the following equation

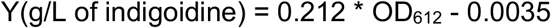

The purity of indigoidine samples were confirmed using H-NMR as previously described^36^. Indigoidine yields were calculated assuming complete utilization of glucose or *p*CA based on the amount of fed substrate in minimal media containing no other carbon sources.

### Advanced micro bioreactor method: 250 mL ambr^®^ 250 bioreactor cultivations

Fed-batch bioreactor experiments were carried out in a 12-way ambr^®^ 250 bioreactor system equipped with 250 mL single-use, disposable bioreactors (microbial vessel type). The vessels were initially filled with 150 mL M9 minimal salt media (NREL formulation) containing 10 g/L glucose or 8.2 g/L *p*CA as carbon source. Temperature was maintained at 30 °C throughout the fermentation process and agitation was set constant to 1300 rpm. Airflow was set constant to 0.5 VVM based on the initial working volume and pH was maintained at 7.0 using 4 N NaOH. Reactors were inoculated manually with 5 mL of pre-culture cell suspension for an initial OD_600_ of ~0.1. After an initial batch phase of 12 hours, cultures were fed with a concentrated feed solution (86 g/L *p*CA, 120 g/L ammonium sulfate, 50 μg/mL kanamycin, 20 mM arabinose) by administering feed boluses every two hours restoring *p*CA concentrations to 8.2 g/L (feed parameters: 3.1 min @ 50 mL/h). Samples were taken 1-2 times every day (2 mL volume) and stored at −20 °C. The ambr^®^ 250 runtime software and integrated liquid handler was used to execute all process steps.

### RbTnSeq fitness experiment under different bioreactor conditions

RbTnSeq fitness assay/experiment was carried out as previously reported^22,25^. Briefly, pooled *P. putida* KT2440 pooled transposon libraries were thawed from 500 μL glycerol stocks and inoculated into 25 mL LB media. Cultures were grown overnight to saturation at 30 °C with shaking (200 rpm). The culture was then subcultured three times in M9 minimal salt media to prepare the seed culture used for bioreactor inoculation. Each of the bioreactors were inoculated to a starting optical density 600 nm at ~0.2. A 2 L bioreactor equipped with a Sartorius BIOSTAT B^®^ fermentation controller (Sartorius Stedim Biotech GmbH, Goettingen, Germany), fitted with two Rushton impellers fixed at an agitation speed of 800 rpm was used. The temperature was held constant at 30 °C. The bioreactor pH was monitored using the Hamilton EasyFerm Plus PHI VP 225 Pt100 (Hamilton Company, Reno, NV) and was maintained at a pH of 7 using 10 M sodium hydroxide. Dissolved oxygen concentration was monitored using Hamilton VisiFerm DO ECS 225 H0. Initial reactor volume was 1 L M9 Minimal Media (10 g/L glucose, 2 mM magnesium sulfate, 0.1 mM calcium chloride, 12.8 g/L sodium phosphate dibasic heptahydrate, 3 g/L potassium phosphate monobasic, 0.5 g/L sodium chloride and 1 g/L ammonium chloride), and 50 mL overnight pre-culture in the same media. For fed-batch experiments, the feeding solution contained 100 g/L glucose, and 300 mM ammonium chloride. The dissolved oxygen (DO) was maintained at either 10% DO or 30% DO in respective bioreactors. The feed rate was set at 1 g/hr glucose and 3 mM NH4Cl with at 10% or 30% DO as indicated. 1 mL samples were harvested, and a cell pellet was collected by centrifugation. Refer to Table 1 for a full description of parameters used in each experiment. As needed, a 1 mL bolus of anti-foam B (Sigma Aldrich) was injected into the bioreactor to control excessive foam formation. Several bioreactor runs were excluded from this analysis if the barcode diversity in the RB-TnSeq data pipeline failed quality check steps. Genomic DNA was extracted and processed for library generation and barcode quantification by Illumina sequencing as previously described^22^. The fitness data described in this work will be available upon publication at http://fit.genomics.lbl.gov.

To assess the statistical significance of each fitness value, a *t*-like test statistic (t-score) of the form fitness/sqrt (estimated variance) was used as described previously in Wetmore *et al^22^*. A gene was considered to have an enhanced fitness phenotype in an experiment if fitness >1.5, *t* > 2 and have a fitness defect when fitness value was < −2, *t* < −2 (and |fitness| > 95th percentile(|fitness|) + 0.5, as described previously^54^). Hierarchical clustering and heatmap visualization in Figure 1 and 2 were done using Python library Seaborn 0.11.1^55^.

### Constraint Based modeling to select metabolic gene deletion strains

*Pseudomonas putida* KT2440 genome scale metabolic model (GSM) iJN1462^35^ was modified to account for indigoidine biosynthesis and used to identify a gene knockout strategy that impacted indigoidine flux. Aerobic growth with either glucose or *para*-coumarate (*p*CA) as the sole carbon source was used to model growth. The ATP maintenance demand was kept the same (0.97 mmol/gDW/h) whereas glucose uptake rate and *p*CA uptake rate were set at 6.3 mmol/gDW/h^56^ and 4.04 mmol/gDW/h^57^ respectively. Flux Balance Analysis (FBA) was used to calculate the maximum theoretical yield (MTY) from reaction stoichiometry and redox balance and also for single gene deletion analysis. Minimization of metabolic adjustment (MOMA) analysis^28^ was used to predict single gene deletion with minimum perturbation in the metabolic flux distribution compared to wild type. Flux variability analysis (FVA) was used to check for minimum and maximum indigoidine flux for each gene deletion strain. COBRA Toolbox v.3.0^58^ in MATLAB R2017b was used for FBA, FVA and MOMA simulations with the GLPK (https://gnu.org/software/glpk) or Gurobi optimization solvers.

## List of Supplementary Tables, Figures, and Datasets

**Supplementary Table 1**: Details for candidate deletion strains identified from Rb-TnSeq mutant library in bioreactor cultivation with enhanced fitness. Each precise deletion contains a common ~250bp sequence of DNA derived from the budding yeast *SMC1* gene and a unique 10bp DNA sequence to aid in identification.

**Supplementary Table 2:** Genome-scale metabolic model derived maximum theoretical yield of alpha-ketoglutarate, glutamine and indigoidine from glucose or *para*-coumarate (*p*CA) with respect to stoichiometry and redox balance in *P. putida.*

**Supplementary Table 3:** Evaluation of gene deletion targets with enhanced fitness from RB-TnSeq profiling for impact on indigoidine production

**Supplementary Table 4:** Fitness profile of PP_4099 mutant across other conditions in the RB-TnSeq fitness browser. Refer to Figure 4A.

**Supplementary Table 5:** Bioinformatic analysis of potential GacA regulated genes in *P. putida* compared to the *P. aeruginosa* regulatory network for GacA as described by Huang *et al,* 2019^43^.

**Supplementary Table 6:** List of plasmids used in this study.

**Supplementary Table 7:** List of strains used in this study.

**Supplementary Figure 1.** Metabolic pathway showing utilization of glucose or the lignin derived aromatic *para*-coumarate (*p*CA) for the production of heterologous bioproduct indigoidine. Indigoidine is derived from the TCA intermediate alpha-ketoglutarate (AKG) via two molecules of glutamine. Adapted from Johnson *et al,* 2019^12^.

**Supplementary Figure 2:** Indigoidine peoduciton in ΔPP_1109 deletion strains. Production of indigoidine from a genomically integrated pathway was conducted as described in Figure 5.

**Supplementary Figure 3:** String database connectivity map of PP_4099/*gacA*. Genes represented on the left connectivity map by their respective gene names are PP_0401/*ksgA,* PP_1623/*rpoS,* PP_1650/*gacS,* PP_1656/*relA,* PP_4097/*pgsA*, PP_4098/*uvrC* and PP_4099/*gacA*. Lower left subnetwork in the right connectivity map represents genes involved in glutamate/glutamine biosynthesis. Genes represented by their respective gene names are PP_0675/*gdhA*, PP_5075/*gltD* and PP_5076/*gltB.*

**Supplementary Figure 4.** A Systems Biology Approach to Characterize Determinants of Bioreactor Fitness. A. Workflow to identify and build new platform strains with increased bioreactor fitness using transposon mutant library. B. We used our mutant library to study the efficient bioconversion of lignin derived aromatic monomer, *para*-coumarate, into a higher value product, a renewable pigment, indigoidine. A representative *P. putida* clone expressing the heterologous indigoidine pathway is shown. C. Strain productivity was characterized at both laboratory and industrially relevant scales.

**Supplementary Data 1:** RBTnSeq gene mutants with decreased fitness.

**Supplementary Data 2:** RBTnSeq gene mutants with increased fitness.

## Data availability

Data supporting the findings of this work are available within the paper and its supplementary information files. The fitness data described in this work will be available upon publication at http://fit.genomics.lbl.gov. A reporting summary for this article is available as a supplementary information file. List of plasmids used in this study are described in **Supplementary Table 6** and their sequences are available at public-registry.jbei.org (registration of a free account is required). All strains used in this study are described in **Supplementary Table 7** and may be also requested from public-registry.jbei.org. Additional requests for datasets and strains generated and analyzed during the current study are available from the corresponding author upon request.

## Acknowledgements

We thank Christopher J. Petzold, Ashish Misra, Megan Garber, Alex Codik, and members of the Mukhopadhyay group for constructive feedback and technical assistance on this work. We also thank Jim Colton (Graphpad Software, San Diego, CA) for assistance using Prism Graphpad for data visualization. The work conducted by the U.S. Department of Energy Joint Genome Institute, a DOE Office of Science User Facility, is supported by the Office of Science of the U.S. Department of Energy under Contract No. DE-AC02-05CH11231.

## Author contributions

Conceptualization of the project: AM, TE, DB. Strain construction, molecular biology, indigoidine quantification: TE, AL, RH, EB and JT. Contributed critical reagents: TE, RH. Interpreted results: AM, TE, AL, RH, DB, AD. RbTnSeq fitness experiment using bioreactors: TE, AL, RH. Ambr250 Fed-Batch Production: AL, JPP, TE and DT. Implementation of Computational Methods: DB. Drafted the manuscript: TE, DB, AM. Raised funds: AM and DT. All authors contributed to and provided feedback on the manuscript and approved the final manuscript.

