## Supplementary Materials for "High Throughput Fitness Profiling Reveals Loss Of GacS-GacA Regulation Improves Indigoidine Production In *Pseudomonas putida*"

**Supplementary Table 1**: Details for candidate deletion strains identified from Rb-TnSeq mutant library in bioreactor cultivation with enhanced fitness. Each precise deletion contains a common ~250bp sequence of DNA derived from the budding yeast *SMC1* gene and a unique 10bp DNA sequence to aid in identification.

| Locus ID | Gene Name | Proposed Function | Allelic Exchange Plasmid Number | Unique DNA Barcode (5'-3') | Gene Essentiality* |
|---|---|---|---|---|---|
| PP_0063 | | putative Lipid A biosynthesis lauroyl acyltransferase | pTE284 | ACGGGCCTTT | |
| PP_0290 | *hisH* | imidazole glycerol phosphate synthase subunit HisH | pTE255 | AACCCCAAGA | essential |
| PP_0393 | | putative 2-amino-4-hydroxy-6-hydroxymethyldihydropteridine pyrophosphokinase | pTE287 | TTTCCGGAAT | essential |
| PP_0574 | *rsbU* | LuxR family transcriptional regulator | pTE272 | CAGCGATTTA | |
| PP_0856 | *yfgL* | lipoprotein | pTE262 | TAACCTCCCA | |
| PP_0887 | *prrB/ regB* | sensor histidine kinase | pTE257 | AGGCCGTTC | |
| PP_0888 | *regA* | photosynthetic apparatus regulatory protein RegA | pTE256 | TCCGGCCGGT | |
| PP_1109 | | GntR family transcriptional regulator | pTE270 | TGCTGATTTC | |
| PP_1111 | | putative Ser-tRNA(Ala) deacylase; Gly-tRNA(Ala) deacylase | pTE269 | GCGGACAATT | essential |
| PP_1216 | *ruvA* | Holliday junction ATP-dependent DNA helicase RuvA | pTE263 | GACAGACAAA | essential |
| PP_1217 | *ruvB* | Holliday junction ATP-dependent DNA helicase RuvB | pTE264 | AATTTCTGTG | |
| PP_1233 | *tusA-I* | sulfurtransferase | pTE258 | CTCGCCTCCT | essential |

| | | | | | |
|---|---|---|---|---|---|
| PP_1385 | *ttg1B* | RND efflux pump, toluene responsive | pTE289 | TACTGGCACC | |
| PP_1428 | *mucA* | sigma factor AlgU negative regulator | pTE254 | TAACAACAGT | essential |
| PP_1656 | | ATP:GTP 3'-pyrophosphotransferase | pTE286 | TGGTAATCCC | |
| PP_1733 | *minD* | MinC-MinD-MinE system ATPase | pTE266 | CATGGCCTTA | essential |
| PP_1734 | *minC* | septum site-determining protein MinC | pTE267 | AGCGCGCATT | |
| PP_2336 | | aconitate hydratase 1 | pTE288 | TACCCTGGAC | |
| PP_2889 | | transmembrane regulator PrtR | pTE261 | TGGGATGGCT | |
| PP_3052 | | hypothetical protein | pTE253 | TGCCGAATCA | essential |
| PP_4099 | *gacA/ uvrY* | GacA/GacS (BarA/UvrY) two-component system response regulator | pTE232 | TGGGCTTGCG | |
| PP_4120 | *nuoB* | NADH-quinone oxidoreductase subunit B | pTE278 | AATAGGACCA | |
| PP_4121 | *nuoC* | NADH-quinone oxidoreductase subunit C/D | pTE279 | GGTTAATTGG | |
| PP_4124 | *nuoG* | NADH-quinone oxidoreductase subunit G | pTE280 | CTTCTTCAAA | |
| PP_4129 | *nuoL* | NADH:ubiquinone oxidoreductase, membrane subunit L | pTE281 | GAAGTTACCC | |
| PP_4131 | *nuoN* | NADH-quinone oxidoreductase subunit N | pTE282 | TACCTAGGAC | essential |
| PP_4799 | | muramoyltetrapeptide carboxypeptidase | pTE259 | TTACGTTCCA | essential |
| PP_4804 | *rlpA* | RlpA-like lipoprotein | pTE260 | AATTAATTAA | essential |
| PP_5146 | *rppH* | RNA pyrophosphohydrolase | pTE268 | TTCTTCTGGG | essential |
| PP_5227 | *lysA-II* | diaminopimelate decarboxylase | pTE283 | AGAGAATCAA | |

| | | | | |
|---|---|---|---|---|
| PP_5309 | *oxyR* | oxidative and nitrosative stress transcriptional dual regulator | pTE271 | ACACGTCACA |
| PP_5310 | *recG* | junction-specific ATP-dependent DNA helicase | pTE265 | GTGAACTGGC |
| PP_5338 | *aspA* | aspartate ammonia-lyase | pTE285 | ATATATTAGG |

*Genes for which a deletion could not be recovered by *sacB* counterselection on LB agar plates supplemented with 10% (w/v) sucrose are presumed to be essential. For each locus tested, at least 2 independent single-crossover clones were generated and at least 100 kanS sucR clones were genotyped.

**Supplementary Table 2:** Genome-scale metabolic model derived maximum theoretical yield of alpha-ketoglutarate, glutamine and indigoidine from glucose or *para*-coumarate (*p*CA) with respect to stoichiometry and redox balance in *P. putida.*

| Metabolite | mol/mol of glucose | mol/mol of *p*CA |
|---|---|---|
| Alpha-ketoglutarate | 1.329 | 1.651 |
| Glutamine | 1.148 | 1.408 |
| Indigoidine | 0.541 | 0.660 |

**Supplementary Table 3:** Evaluation of gene deletion targets with enhanced fitness from RB-TnSeq profiling for impact on indigoidine production.

| Gene | Gene Name | Associated Reactions in iJN1462[1] | Biomass (gDCW/ mmol of glucose) | Indigoidine MTY (mol/mol of glucose) | Biomass (gDCW/ mmol of *p*CA) | Indigoidine MTY (mol/mol of pCA) |
|---|---|---|---|---|---|---|
| WT | - | - | 0.1 | 0.54 | 0.11 | 0.66 |
| PP_4124 | *nuoG* | NADH16pp | 0.04 | 0.48* | 0.05 | 0.57 |
| PP_4131 | *nuoN* | NADH16pp | 0.04 | 0.48* | 0.05 | 0.57 |
| PP_4943 | | GTR2_kt, LPSGTR2_kt | 0.1 | 0.54 | 0.11 | 0.66 |
| PP_5227 | *lysA-2* | DAPDC | 0.1 | 0.54 | 0.11 | 0.66 |
| PP_0063 | *htrB* | EDTXS2_kt, EDTXS5_kt, EDTXS1_kt, EDTXS6_kt | 0.1 | 0.54 | 0.11 | 0.66 |
| PP_5338 | *aspA* | ASPT | 0.1 | 0.54 | 0.11 | 0.66 |
| PP_1656 | *relA* | GDPDPK, GTPDPK | 0.1 | 0.54* | 0.11 | 0.66 |
| PP_0290 | *hisH* | IG3PS | 0 | 0.54 | 0 | 0.66 |
| PP_0393 | *folK-1* | HPPK | 0.1 | 0.54 | 0.11 | 0.66 |
| PP_2336 | *acnM* | MICITDr | 0.1 | 0.54 | 0.11 | 0.66 |
| PP_4129 | *nuoL* | NADH16pp | 0.04 | 0.48* | 0.05 | 0.57 |
| PP_4799 | | 4PCP, AM4PCP, UM4PCP, AGM4PCP | 0.1 | 0.54 | 0.11 | 0.66 |
| PP_4120 | *nuoB* | NADH16pp | 0.04 | 0.48* | 0.05 | 0.57 |
| PP_4121 | *nuoCD* | NADH16pp | 0.04 | 0.48* | 0.05 | 0.57 |
| PP_1385 | *ttgB* | APCt5, APCt6, CMt2ex, CMtpp, INDOLEt5, INDOLEt6, MXYLt5, MXYLt6, OXYLt5, OXYLt6, PXYLt5, PXYLt6, TOLt5, TOLt6, TTRCYCt5, TTRCYCtpp | 0.1 | 0.54 | 0.11 | 0.66 |

*FBA solution as no solution exists when using MOMA analysis

**Supplementary Table 4:** Fitness profile of PP_4099 mutant across other conditions in the RB-TNSeq fitness browser. Refer to Figure 4A.

| Condition* | PP_4099 | PP_0063 | PP_5388 |
|---|---|---|---|
| 1,2-Propanediol (C) | 1.1 | 0.1 | 0 |
| 1,2-Propanediol (C) | 1.1 | 0.3 | -0.2 |
| 1,3-Butandiol (C) | 1.7 | -0.3 | 0 |
| 1,3-Butandiol (C) | 0.5 | -0.2 | -0.1 |
| 1,4-Butanediol (C) | 0.1 | 0.2 | 0.1 |
| 1,4-Butanediol (C) | -0.9 | -0.4 | 0 |
| 1,5-Pentanediol (C) | 3.5 | -0.6 | 0 |
| 1,5-Pentanediol (C) | 3.7 | 0 | 0.1 |
| 1-Pentanol (C) | 1.4 | -0.3 | 0 |
| 1-Pentanol (C) | 1.3 | -0.7 | -0.1 |
| 2-methyl-1-butanol (C) | 2 | -0.3 | 0.2 |
| 2-methyl-1-butanol (C) | 2 | -0.8 | 0.2 |
| Valerolactam (C) | 1.1 | -0.5 | 0.1 |
| Valerolactam 10 mM (C) | 3.9 | 0.4 | 0 |
| 3-methyl-1-butanol (C) | 1.6 | -2.8 | 0.1 |
| 3-methyl-1-butanol (C) | 2.5 | -0.7 | 0 |
| 3-methyl-3-butenol (C) | 1.6 | -0.4 | -0.2 |
| 3-methyl-3-butenol (C) | 2.2 | 0 | -0.2 |

| | | | |
|---|---|---|---|
| 4-Aminobutyric 10 mM (C) | 3.2 | -0.3 | -0.4 |
| 4-Hydroxybenzoic Acid (C) | 2.4 | -0.8 | 0.1 |
| 4-Hydroxybenzoic Acid (C) | 2 | -0.4 | 0.1 |
| 4-Hydroxyvalerate (C) (40mM) | 3.2 | 0.1 | 0.1 |
| 4-Hydroxyvalerate (C) (40mM) | 4.2 | 0 | -0.1 |
| 5-Aminovaleric 10 mM (C) | 2 | -1.3 | -0.2 |
| 5-Aminovaleric 10 mM (C) | 2.4 | 0 | -0.1 |
| benzoic (C) | 1.2 | -0.8 | -0.2 |
| benzoic (C) | 1.8 | -1.2 | -0.1 |
| benzoic (C) | 1.3 | -0.6 | -0.3 |
| Butanol (C) | 0.7 | -0.6 | 0.1 |
| Butanol (C) | 0.6 | 0.1 | -0.2 |
| Butyl stearate (C) | -0.2 | -1.7 | -0.3 |
| Butyl stearate (C) | 0.3 | -1.8 | 0 |
| D-Glucose (C) | 1.5 | -0.5 | -0.2 |
| D-Glucose (C) | 1.3 | -0.2 | -0.1 |
| D-Glucose (C) | 1.7 | -0.5 | -0.4 |
| D-Glucose (C) | 0.9 | -0.2 | -0.1 |
| D-Glucose (C) | 0.9 | -0.6 | -0.1 |
| Glucose (C) (20mM) | 1 | -0.5 | -0.3 |
| Glucose (C) (20mM) | 1 | -0.3 | 0.2 |

| | | | |
|---|---|---|---|
| Glucose (C) (40mM) | 1.3 | -0.2 | -0.2 |
| Glucose (C) (40mM) | 1.3 | -0.7 | 0.4 |
| D-Lysine 10 mM (C) | 2.5 | -0.9 | 0 |
| D-Lysine 10 mM (C) | 3.2 | -1.5 | 0.4 |
| Decanoic (C) | 2.1 | -1 | -0.5 |
| Decanoic (C) | 2.3 | -2.3 | -0.3 |
| Ethanol (C) | 1.1 | 0 | 0.2 |
| Ethanol (C) | 0.8 | -0.1 | -0.1 |
| Ferulic Acid (C) | 5 | 0.1 | -0.2 |
| Ferulic Acid (C) | 3.6 | -0.2 | 0 |
| Ferulic Acid (C) | 3.5 | 0.4 | -0.2 |
| Heptanoic (C) | -0.2 | -2.3 | -0.2 |
| Heptanoic (C) | 0.3 | -2.3 | -0.3 |
| Hexanoic (C) | 4.6 | -1.3 | -0.6 |
| Hexanoic (C) | 2.9 | -1.2 | -0.9 |
| L-Lysine 10 mM (C) | 4.5 | -0.5 | 0.3 |
| Lauric (C) | 1.5 | -1.6 | -0.3 |
| Lauric (C) | 1.5 | -1.8 | 0.2 |
| Levulinic Acid (C) | 3.4 | -0.5 | -0.2 |
| Levulinic Acid (C) | 3.5 | -0.7 | 0 |
| Levulinic Acid (C) | 1.7 | 1 | 0.1 |

| | | | |
|---|---|---|---|
| Levulinic Acid (C) | 2.7 | 1.3 | -0.2 |
| Myristic (C) | 1 | -2.3 | -0.1 |
| Myristic (C) | 1.5 | -2.6 | -0.2 |
| Nonanoic (C) | 1.4 | -2.2 | 0.2 |
| Nonanoic (C) | 2 | -2.6 | -0.3 |
| Octanoic (C) | 2.2 | -0.7 | -0.5 |
| Octanoic (C) | 1.9 | -1.4 | 0.1 |
| Oleic (C) | 0.2 | -2.6 | 0.2 |
| Oleic (C) | -0.3 | -2.1 | 0.2 |
| p-Coumaric (C) | 1.9 | -0.3 | -0.3 |
| p-Coumaric (C) | 1.9 | -0.7 | 0 |
| p-Coumaric (C) | 1.8 | -0.8 | -0.2 |
| p-Coumaric (C) | 1.8 | -1 | 0.1 |
| Phenylacetic (C) | 3.8 | -1.2 | -0.3 |
| Phenylacetic (C) | 3.2 | -0.7 | 0.2 |
| Acetate (20mM) (C) | 4.9 | 0.1 | 0.3 |
| Acetate (20mM) (C) | 5 | -0.2 | 0.1 |
| Acetate (5mM) (C) | 3.8 | -0.1 | 0.2 |
| Acetate (5mM) (C) | 3.7 | 0 | 0.1 |
| Protocatechuic Acid (C) | 3.7 | -0.9 | 0.3 |
| Protocatechuic Acid (C) | 3 | -0.3 | -0.2 |

| | | | |
|---|---|---|---|
| butyrate (C) | 5.4 | -0.7 | -0.6 |
| butyrate (C) | 5.2 | -1.4 | -0.2 |
| propionate (C) | 3 | -0.4 | 0 |
| propionate (C) | 3.1 | 0.1 | 0.1 |
| Tween 20 (C) | 1.4 | -1.3 | -0.9 |
| Tween 20 (C) | 1 | -1.5 | 0.3 |
| Valeric (C) | 3 | -1.3 | 0.3 |
| Valeric (C) | 3.3 | -2.3 | -0.1 |
| Vanillic Acid (C) | 1.3 | -0.5 | 0.1 |
| Vanillic Acid (C) | 1.3 | 0.1 | -0.1 |
| Vanillin (C) | 4 | -1.4 | -0.2 |
| Vanillin (C) | 3.5 | -0.5 | -0.2 |
| Vanillin (C) | -2.4 | -0.1 | -0.1 |
| Vanillin (C) | -2.2 | -0.7 | 0.4 |

*Condition details can be found using the Fitness browser at: https://bit.ly/38oy971

**Supplementary Table 5:** Bioinformatic analysis of potential GacA regulated genes in *P. putida* compared to the *P. aeruginosa* regulatory network for GacA as described in Huang *et al*, 2019[2].

| ***P. aeruginosa* Gene Loci** | **Gene Name** | ***P. putida* Gene Loci** | **% Similarity or Identity** | **RbTnSeq Fitness under bioreactor condition** |
|---|---|---|---|---|
| PA0048 | | PP_4133 | 27% | Negative |
| PA0073 | *tagT1* | PP_0507 | 41% | Negative |
| PA0091 | *vgrG1* | PP_3106 | 36% | Neutral |
| PA0354 | | PP_3386 | 33% | Neutral |
| PA0509 | *nirN* | PP_2669 | 25% | Neutral |
| PA0630 | | PP_5742 | 25% | Neutral |
| PA0688 | *lapA* | PP_2656 | 22% | Neutral |
| PA0746 | *PA0746* | PP_2216 | 46% | Slightly negative |
| PA0751 | *PA0751* | PP_1415$ | 68% | Slightly negative |
| PA0835 | *pta* | PP_0774$ | 78% | Neutral |
| PA0898 | *aruD/astD* | PP_4478$ | 82% | Neutral |
| PA0899 | *aruB* | PP_4477$ | 84% | Neutral |
| PA0903 | *alaS* | PP_4474$ | 86% | No data; Essential gene |
| PA1107 | *roeA* | PP_1411 | 48% | Neutral |
| PA1197 | | PP_5402 | 37% | Neutral |
| PA1663 | *sfa2* | PP_4696 | 45% | Neutral |
| PA1712 | exsB | No homolog | No homolog | No homolog |
| PA1906 | | No homolog | No homolog | No homolog |

| | | | | |
|---|---|---|---|---|
| PA2586 | *gacA* | PP_4099 | | Strong, positive |
| PA2683 | | PP_3191 | 38% | None |
| PA2727 | | PP_3691 | 32% | Neutral |
| PA2729 | | No homolog | No homolog | No homolog |
| PA3309 | | PP_2132 | 29% | Slightly negative |
| PA3328 | | PP_3944$ | 38% | Neutral |
| PA3329 | | PP_4243 | 28% | Neutral |
| PA3330 | | PP_1274 | 36% | Neutral |
| PA3416 | | PP_4402 | 42% | Neutral |
| PA3417 | | PP_4401 | 37% | Neutral |
| PA3457 | | PP_0054$ | 38% | Slightly negative; not in all bioreactor replicates |
| PA3533 | *grxD* | PP_1081$ | 90% | No data; Essential gene |
| PA3560 | *fruA* | PP_0795$ | 73% | Neutral |
| PA3568 | *prpE* | PP_2351$ | 78% | Neutral |
| PA3569 | *mmsB* | PP_4666$ | 53% | Neutral |
| PA3572 | | PP_5598$ | 48% | Neutral |
| PA3637 | *pyrG* | PP_1610$ | 89% | Essential gene |
| PA3727 | | No homolog | No homolog | No homolog |
| PA3728 | | No homolog | No homolog | No homolog |
| PA3873 | *narJ* | No homolog | No homolog | No homolog |

| | | | | |
|---|---|---|---|---|
| PA4112 | *barA/ gacS* | PP_1650 | 38% | Strong, positive |
| PA4218 | *ampP* | PP_1355 | 33% | Slightly negative |
| PA4219 | *ampO* | PP_0349 | 27% | Inconsistent results |
| PA4221 | *fptA* | PP_4217 | 35% | Negative |
| PA4224 | *pchG* | No homolog | No homolog | No homolog |
| PA4225 | *pchF* | PP_4219 | 29% | Neutral |
| PA4438 | | PP_1312[$] | 81% | No data; Essential gene |
| PA4488 | *magE* | No homolog | No homolog | No homolog |
| PA4587 | *ccpR* | PP_2943[$] | 41% | Neutral |
| PA4611 | | No homolog | No homolog | No homolog |
| PA4676 | | PP_0100 | 34% | No data; Essential gene |
| PA4705 | *phuW* | PP_4686[$] | 56% | Slightly negative |
| PA4925 | | PP_1353 | 33% | Positive |
| PA5113 | | No homolog | No homolog | No homolog |
| PA5475 | | PP_3232[$] | 58% | Neutral |
| PA5531 | *tonB1* | PP_4994 | 35% | Negative in all bioreactors |
| PA0527.1 | *rsmY* | Near PP_0370 [3] | | Not annotated; No data |
| PA3621.1 | *rsmZ* | Near PP_1624 [3] | | Not annotated; No data |

| | | | | |
|---|---|---|---|---|
| PA0592# | *ksgA/ rrnaAD* | PP_0401$ | 79% | neutral |
| PA0600# | | PP_0409$ | 65% | Slightly negative |
| PA3622# | *rpoS* | PP_1623$ | 87% | Negative |
| PA0934# | *relA* | PP_1656$ | 87% | Positive |
| PA0600# | | PP_2127 | 35% | Neutral |
| PA2584# | *pgsA* | PP_4097$ | 84% | No data; Essential gene |
| PA2585# | *uvrC* | PP_4098$ | 84% | Positive |
| PA1898# | *qscR/ppoR* | PP_4647$ | 29% | Neutral |

$ - ortholog according to Biocyc database

### - candidate genes from a StringDB analysis

**Supplementary Table 6:** List of plasmids used in this study.

| JBEI Accession ID | Plasmid Number | Genotype | Reference[1,2] |
|---|---|---|---|
| JBEI-145804 | pTE232 | *pEX18GM-ΔPP_4099 aacC1 sacB* | This study |
| JBEI-145806 | pTE253 | *pEX18GM-ΔPP_3052 aacC1 sacB* | This study |
| JBEI-145808 | pTE254 | *pEX18GM-ΔPP_1428 aacC1 sacB* | This study |
| JBEI-145810 | pTE255 | *pEX18GM-ΔPP_0290 aacC1 sacB* | This study |
| JBEI-145812 | pTE256 | *pK18mobsacB-ΔPP_0888 kanR sacB* | This study |
| JBEI-145814 | pTE257 | *pEX18GM-ΔPP_0887 aacC1 sacB* | This study |
| JBEI-145816 | pTE258 | *pEX18GM-ΔPP_1233 aacC1 sacB* | This study |
| JBEI-145818 | pTE259 | *pEX18GM-ΔPP_4799 aacC1 sacB* | This study |
| JBEI-145820 | pTE260 | *pK18mobsacB-ΔPP_4804 kanR sacB* | This study |
| JBEI-145822 | pTE261 | *pEX18GM-ΔPP_2889 aacC1 sacB* | This study |
| JBEI-145824 | pTE262 | *pEX18GM-ΔPP_0856 aacC1 sacB* | This study |

| | | | |
|---|---|---|---|
| JBEI-145826 | pTE263 | *pEX18GM-ΔPP_1216 kanR sacB* | This study |
| JBEI-145828 | pTE264 | *pEX18GM-ΔPP_1217 aacC1 sacB* | This study |
| JBEI-145830 | pTE265 | *pEX18GM-ΔPP_5310 aacC1 sacB* | This study |
| JBEI-145832 | pTE266 | *pEX18GM-ΔPP_1733 aacC1 sacB* | This study |
| JBEI-145834 | pTE267 | *pK18mobsacB-ΔPP_1734 kanR sacB* | This study |
| JBEI-145836 | pTE268 | *pEX18GM-ΔPP_5146 aacC1 sacB* | This study |
| JBEI-145838 | pTE269 | *pEX18GM-ΔPP_1111 aacC1 sacB* | This study |
| JBEI-145840 | pTE270 | *pEX18GM-ΔPP_1109 aacC1 sacB* | This study |
| JBEI-145842 | pTE271 | *pEX18GM-ΔPP_5309 aacC1 sacB* | This study |
| JBEI-145844 | pTE272 | *pEX18GM-ΔPP_0574 aacC1 sacB* | This study |
| JBEI-145846 | pTE278 | *pK18mobsacB-ΔPP_4120 kanR sacB* | This study |
| JBEI-145848 | pTE279 | *pK18mobsacB-ΔPP_4121 kanR sacB* | This study |
| JBEI-145850 | pTE280 | *pK18mobsacB-ΔPP_4124 kanR sacB* | This study |

| | | | |
|---|---|---|---|
| JBEI-145852 | pTE281 | *pK18mobsacB-ΔPP_4129 kanR sacB* | This study |
| JBEI-145854 | pTE282 | *pK18mobsacB-ΔPP_4131 kanR sacB* | This study |
| JBEI-145856 | pTE283 | *pK18mobsacB-ΔPP_5227 kanR sacB* | This study |
| JBEI-145858 | pTE284 | *pK18mobsacB-ΔPP_2889 kanR sacB* | This study |
| JBEI-145860 | pTE285 | *pK18mobsacB-ΔPP_5338 kanR sacB* | This study |
| JBEI-145862 | pTE286 | *pK18mobsacB-ΔPP_1656 kanR sacB* | This study |
| JBEI-145864 | pTE287 | *pK18mobsacB-ΔPP_0393 kanR sacB* | This study |
| JBEI-145866 | pTE288 | *pK18mobsacB-ΔPP_2336 kanR sacB* | This study |
| JBEI-108734 | pTE289 | *pK18mobsacB-ΔPP_1385 kanR sacB* | Mohamed *et al*, 2020[4] |

[1] The vector pEX18GM is described in Hoang *et al* 1998[5].

[2] The vector pK18mobsacB is described in Schäfer *et al* 1994[6].

**Supplementary Table 7:** List of strains used in this study.

| JBEI Accession ID | Strain Number | Genotype | Reference |
|---|---|---|---|
| JBEI-13809 | TEAM-911 | *P. putida* mt-2 KT2440 cmR kanS gntS sucR | ATCC-47054 |
| None[1] | KT.RB-TnSeq | *P. putida* KT2440 RB-TnSeq pooled mutant library kanR | Price *et al,* 2019[7] |
| JBEI-137184 | TEAM-1120 | *P. putida* KT2440 *PP_5402::araC-BADp-bpsA-sfp* | Banerjee *et al,* 2020[8] |
| JBEI-147142 | TEAM-1138 | *P. putida* TEAM-1120 Δ*PP_4099* (ΔgacA) | This study |
| JBEI-147143 | TEAM-1285 | *P. putida PP_5402::araC-BADp-bpsA-sfp ΔPP_4120* kanR | This study |
| JBEI-147144 | TEAM-1382 | *P. putida* TEAM-1120 Δ*PP_4124* | This study |
| JBEI-147146 | TEAM-1291 | *P. putida PP_5402::araC-BADp-bpsA-sfp* Δ*PP_4129* kanR | This study |
| JBEI-147147 | TEAM-1287 | *P. putida PP_5402::araC-BADp-bpsA-sfp* Δ*PP_4121* kanR | This study |
| JBEI-147148 | TEAM-1297 | *P. putida PP_5402::araC-BADp-bpsA-sfp* Δ*PP_5338* kanR | This study |

| | | | |
|---|---|---|---|
| JBEI-147149 | TEAM-1295 | *P. putida PP_5402::araC-BADp-bpsA-sfp ΔPP_0063* kanR | This study |
| JBEI-147150 | TEAM-1293 | *P. putida PP_5402::araC-BADp-bpsA-sfp ΔPP_5227* kanR | This study |
| JBEI-147151 | TEAM-1299 | *P. putida PP_5402::araC-BADp-bpsA-sfp ΔPP_1656* kanR | This study |
| JBEI-147152 | TEAM-1301 | *P. putida PP_5402::araC-BADp-bpsA-sfp ΔPP_2336* kanR | This study |
| JBEI-147153 | TEAM-1303 | *P. putida PP_5402::araC-BADp-bpsA-sfp ΔPP_1385* (*Δttg1B*) kanR | This study |
| JBEI-147154 | TEAM-1495 | *P. putida* TEAM-1120 *ΔPP_1109* | This study |
| JBEI-147155 | TEAM-1493 | *P. putida* TEAM-1120 *ΔPP_2889* | This study |

[1]The *P. putida* RB-TnSeq library is stored at -80 ℃ in 1 mL single use aliquots, which is incompatible with the JBEI strain archiving format.

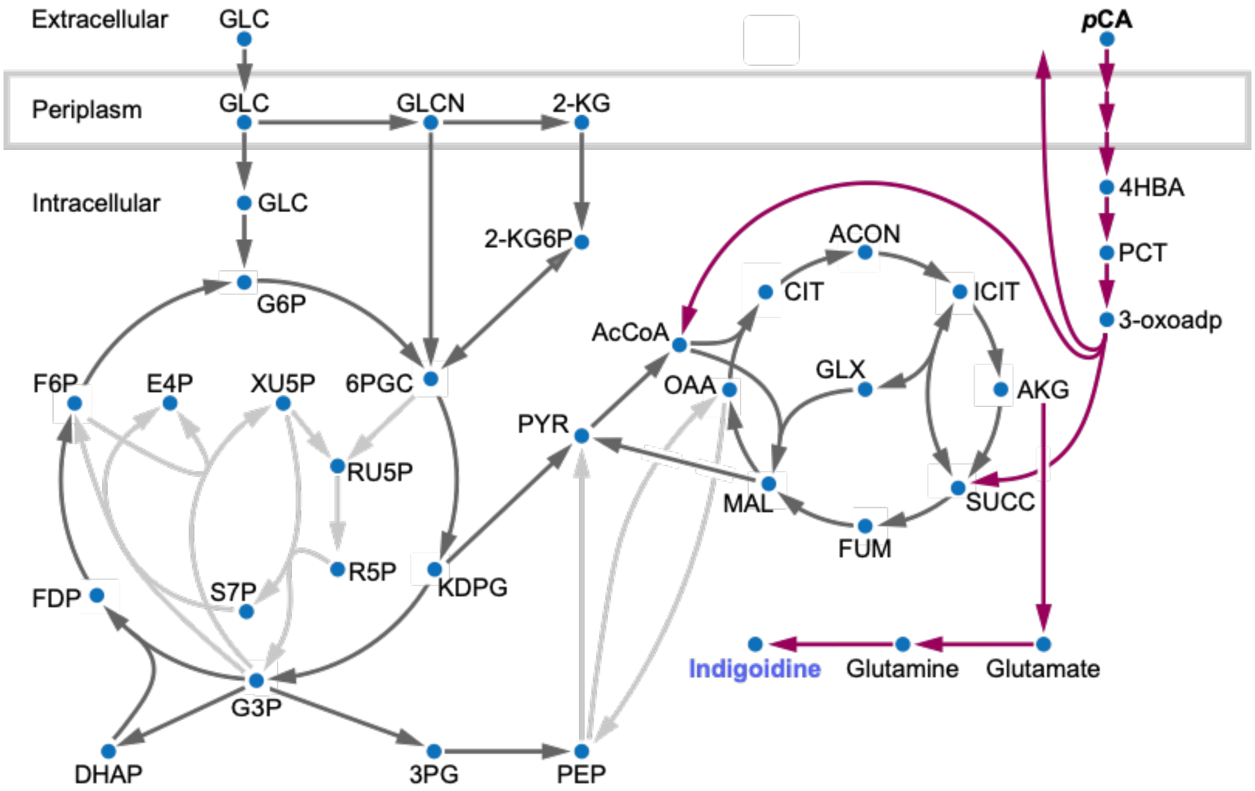

**Supplementary Figure 1.** Metabolic pathway showing utilization of glucose (GLC) or the lignin derived aromatic para-coumarate (*p*CA) for the production of heterologous bioproduct indigoidine. Indigoidine is derived from the TCA intermediate alpha-ketoglutarate (AKG) via two molecules of glutamine. The metabolic flux template was derived from Johnson *et al,* 2019[9]. Other abbreviations include glucose-6-phosphate (G6P), gluconate (GLCN), 2-ketogluconate (2-KG), 2-ketogluconate-6-phosphate (2-KG6P), fructose-6-phosphate (F6P), fructose-1,6-biphosphate (FDP), 6-phosphogluconate (6PGC), 2-keto-3-deoxy-6-phosphogluconate (KDPG), ribulose-5-phosphate (RU5P), ribose-5-phosphate (R5P), xylulose-5-phosphate (XU5P), sedoheptulose-7-phosphate (S7P), erythrose-4-phosphate (E4P), glyceraldehyde-3-phosphate (G3P), glycerate-3-phosphate (3PG), dihydroxyacetone phosphate (DHAP), phosphoenolpyruvate (PEP), pyruvate (PYR), oxaloacetate (OAA), acetyl Coenzyme A (AcCoA), aconitate (ACON), citrate (CIT), isocitrate (ICIT), succinate (SUCC), fumarate (FUM), malate (MAL), glyoxylate (GLX), 4-hydroxybenzoate (4HBA), protocatechuate (PCT) and 3-oxoadipate or beta-ketoadipate (3-oxoadp). Arrows indicate individual metabolic reactions except for the conversion of PCT to 3-oxoadp, which is catalyzed by four enzymatic reactions.

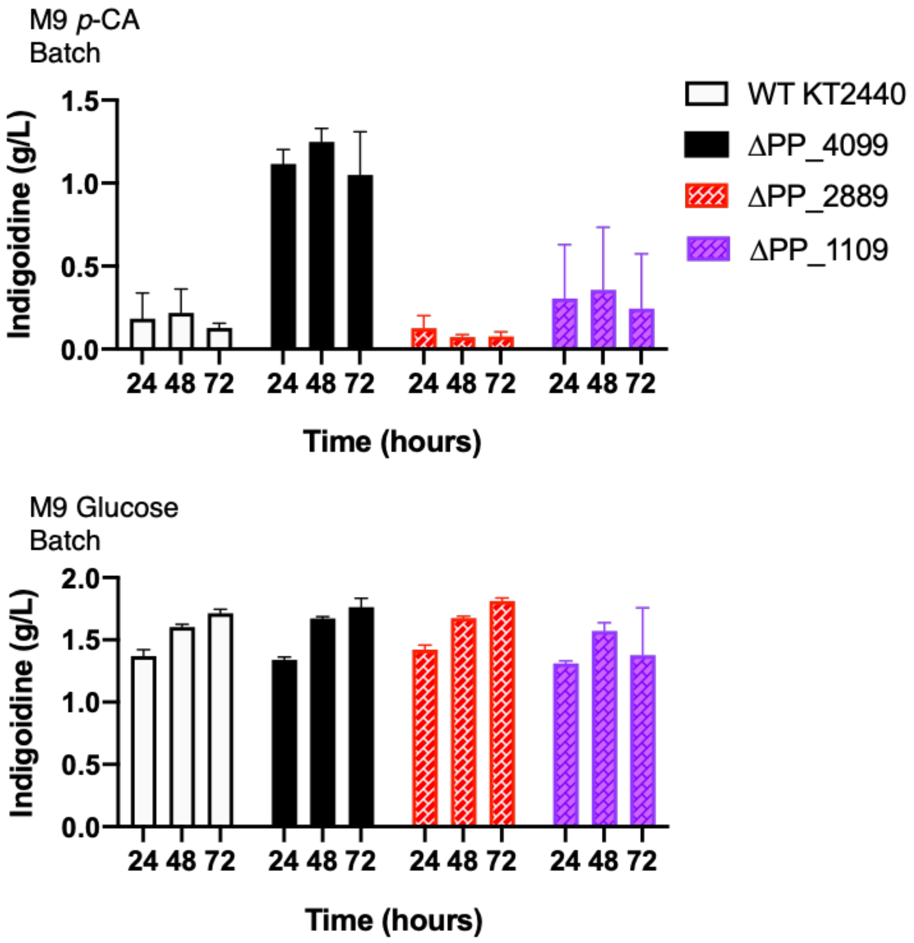

**Supplementary Figure 2:** Indigoidine production in ΔPP_2889 and ΔPP_1109 deletion strains. Production of indigoidine from a genomically integrated pathway was conducted as described in Figure 5. Data are presented as mean ± SD from n = 4 independent biological replicates.

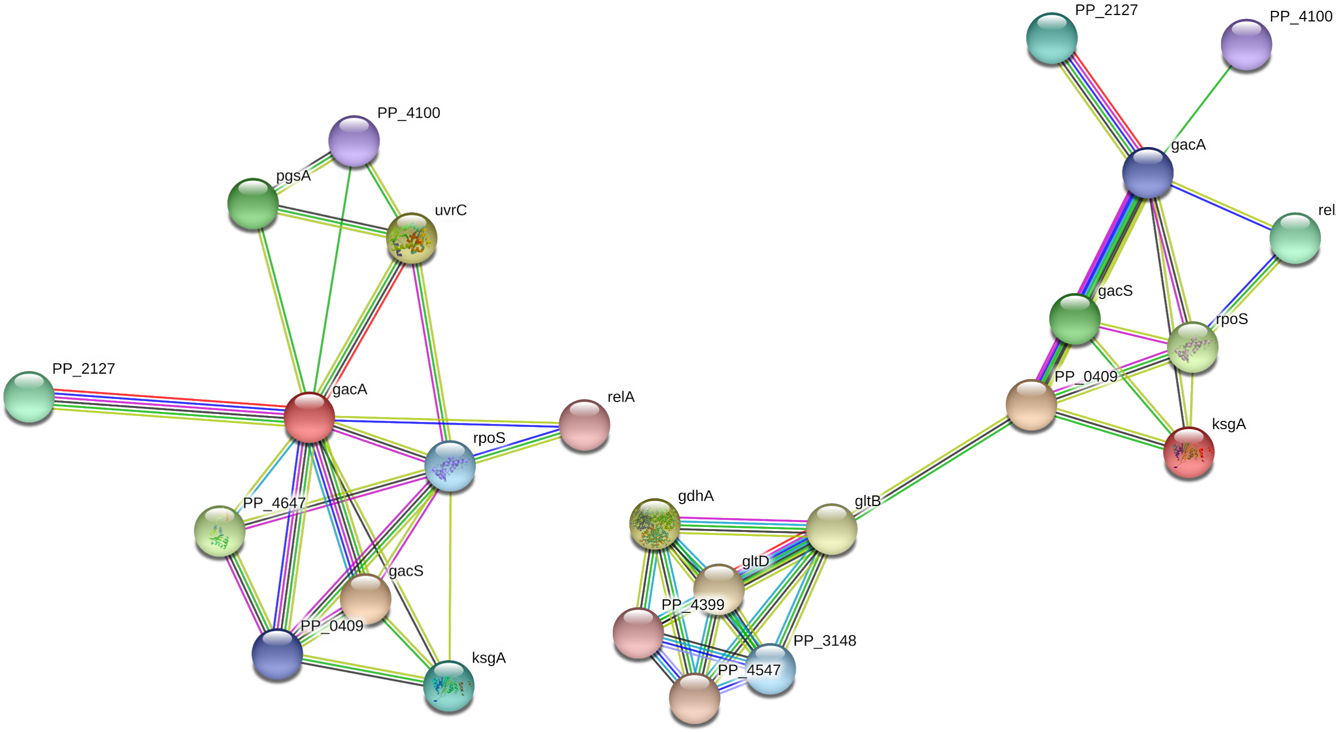

**Supplementary Figure 3:** String database[10] connectivity map of PP_4099/*gacA*. Genes represented on the left connectivity map by their respective gene names are PP_0401/*ksgA,* PP_1623/*rpoS,* PP_1650/*gacS,* PP_1656/*relA,* PP_4097/*pgsA*, PP_4098/*uvrC* and PP_4099/*gacA*. Lower left subnetwork in the right connectivity map represents genes involved in glutamate/glutamine biosynthesis. Genes represented by their respective gene names are PP_0675/*gdhA*, PP_5075/*gltD* and PP_5076/*gltB.*
